# Fiber Diameter and Architecture Direct Three-Dimensional Assembly of Pericytes into Spheroids

**DOI:** 10.1101/2022.08.02.502506

**Authors:** Sharan Sharma, Jennifer C. Hill, Julie A. Phillippi, Amrinder S. Nain

## Abstract

Due to their physiological relevance, multicellular 3D spheroids are actively replacing standard 2D monolayer cultures. How spheroids are formed through the assembly of individual cells in natural fibrous environments that include a mix of diameters and architectures in vivo remains unknown. Here, we demonstrate that the spontaneous assembly of human vasa vasorum-derived pericytes in 3D spheroids depends on the fiber diameter and network architecture. A parallel arrangement of suspended fibers of all tested diameters (200, 500, and 800 nm) leads to the formation of spheroids, while on crosshatch networks, spheroid assembly on larger diameters is absent. The design of fibrous networks of a mix of diameters and architectures leads to the patterning of spheroids in desired locations. Fiber remodeling in parallel arrangements serves as force sensors providing mechanical insights into the assembly dynamics of spheroids and subsequent cell sprouting from spheroids. Translocation and merger of spheroids occur predominantly on parallel fiber networks, while on crosshatch networks, a cellular exchange is observed between spheroids connected with remodeled fibers. Rho kinase inhibition by Y27632 and subsequent wash-off leads to spheroid disintegration and reassembly, thus, highlighting the role of cell contractility in the assembly and integrity of 3D spheroids. Overall, using extracellular mimicking fiber networks of varying diameters and architectures, we report new insights into the 3D dynamics of spheroids which may inform pericyte’s role in vasculogenesis, and (patho)physiological angiogenesis

## Introduction

Flat 2D cell culture, a traditional method of studying cell biology is being replaced by 3D culture models and environments as numerous studies have shown fundamental behavioral differences (morphology, migration, proliferation and adhesion, genetic expression) when cells are cultured on 2D versus with 3D environments.^1,2^ 2D substrates generally tend to be more stiff when compared with *in vivo* microenvironment and they lack many of the biophysical and biochemical properties crucial for mimicking cellular function.^3,4^ 3D cell culture systems, on the other hand, capture the functional and architectural characteristics of native tissues by having controllable features directing cell-cell and cell-substrate interactions that represent either physiological or pathological environments.^3,5,6^

One 3D culture system is created through assembly of cell “spheroids”, which are multicellular spherical aggregates of cells.^3,7,8^ Spheroids are considered more representative of cells within the *in vivo* tissue microenvironment than 2D monolayer cultures due to differences observed in mechanical force exertion, biophysical and biochemical signaling, and electrical coupling to influence morphology, motility, proliferation, differentiation, and genetic expression.^3^ Spheroids are also known to maintain the functionality and characteristics similar to tissues for longer duration.^7^ Because of these advantages, spheroids are increasingly being used for applications such as tissue engineering, regenerative medicine, drug development and screening, and single- and multi-organ-on-a-chip devices integrated with microfluidics.^4,7,9^ Tumor cell-derived spheroid models are also popular for studying cell behavior and determining the effect of putative therapeutic drugs.^3–5,9^

Over the years, multiple techniques have been developed to assemble and maintain spheroids. Spheroids can either be created in scaffold-free systems such as cell suspensions, using the hanging drop method, or using anti-adhesion substrates.^7^ They can also be grown in scaffold-based system like natural extracellular matrices (ECM), hydrogels and other polymer matrices.^3,6,7^ Various techniques such as pellet culture, microfluidics, spinner culture, rotating wall vessels, and liquid overlays have also been applied to assemble and grow spheroids.^6,7^

Several pluripotent and adult tissue cell lines such as human mesenchymal stem cells, human umbilical vein endothelial cells, murine embryonic stem cells, dermal fibroblasts, chondrocytes and those from various tumors such as colon, breast, pancreatic, and brain cancer cells can be grown as spheroids.^4,6,10^ Pericytes are mural cells are predominantly found in the microvasculature with crucial roles in vascular development and remodeling.^11–13^ Human brain microvascular pericytes form spheroids when cocultured with other cells such as endothelial cells and astrocytes, which have been used to study cell behavior, tissue engineering, and drug delivery.^14–16^ Members of our team have previously demonstrated that human pericytes derived from the adventitial microvascular network known as the vasa vasorum spontaneously self-assemble into spheroids when cultured on Matrigel substrates.^17^

Recent literature identified the importance of mechanobiology of spheroid assembly^18^, growth, and subsequent interactions with the surrounding microenvironment.^19–24^ Since natural cellular ECM environments are fibrous, we inquired if fiber dimensions and their architecture regulate aggregation of cells into spheroids and affected spheroid size, shape, and dynamics. To answer these questions, we used tunable 3D fiber networks as a platform to study the biophysical influence of the underlying microenvironment on pericyte spheroids. To the best of our knowledge, our study is the first to report a direct correlation between fiber diameter and architecture in the assembly of 3D spheroids. Parallel arrangement of fibers of all tested diameters lead to spheroid assembly, while spheroid assembly on large diameter crosshatches is absent. This provides unique opportunity to pattern spheroids at desired locations through design of fiber networks. Fiber remodeling in parallel arrangements serves as force sensors providing mechanical insights into the assembly dynamics of spheroids and subsequent cell sprouting from spheroids. Using quantitative microscopy, we also captured various dynamic interactions of the spheroids with individual cells, other spheroids, and fibers, further revealing new and curious behaviors of pericytes as spheroids within 3D fibrous microenvironments.

## Materials and Methods

### Fiber network manufacturing and characterization

The suspended fiber networks were manufactured on flat, hollow metal scaffolds using our previously reported non-electrospinning STEP technique, a fiber spinning technique with control over fiber diameter, spacing and architecture.^25^ Polystyrene (MW: 2,500,000 g/mol; CAS No. 1025; Scientific Polymer Products, Ontario, NY, USA) was dissolved in xylene (CAS No. 1330-20-7; Carolina, Burlington, NC). The solution was extruded via a micropipette (Jensen Global, Santa Barbara, CA, USA). Aligned fiber networks were fabricated by depositing ∼2 µm thick ‘*base’* fibers spaced ∼350 µm apart, over which orthogonal layers of ∼200/500/800 nm *‘aligned’* fibers with ∼10 µm spacing were deposited.^26–28^ Crosshatch fiber networks involved two densely spaced (∼10 µm) orthogonally deposited layers of fibers having the same diameter (∼200/500/800 nm^29,30^). The intersections were fused using Tetrahydrofuran vapors to preserve their architecture (Sup. Figure 1(a)). SEM images were taken with JSM-IT500 InTouchScope™ Scanning Electron Microscope (JEOL Ltd., Akishima, Tokyo, Japan) to characterize the fiber networks.

### Cell culture on nanofiber scaffolds

The scaffolds with fiber networks were mounted in 6-well plates (CAS No. P06-1.5H-N; Cellvis, Mountain View, CA), sterilized with 70% ethanol for 10 mins and washed twice with PBS. The fibers and glass were functionalized with 4 µg/mL Fibronectin (CAS No. F1141-1MG; MilliporeSigma, Burlington, MA, USA) incubation at 37°C for 45 mins to facilitate cell adhesion. Human pericytes were isolated as previously described^17^ from the adventitia of ascending aorta, resected during heart transplantation operations with approval from the Institutional Review Board (University of Pittsburgh, Protocol #STUDY20040179) and under an informed consent process. Pericytes were culture-expanded in StemMACS™ MSC Expansion Media (Order No. 130-091-680; Miltenyi Biotec, North Rhine-Westphalia, Germany) and immortalized using a HPV E6/E7 lentiviral system as previously reported.^31^ Pericytes were sub-cultured from passages 70-85 using 0.1% Trypsin-EDTA (CAS No R001100; ATCC, Manassas, VA) and seeded on fiber networks. The cells were allowed to adhere to the fiber for 30-45 min at 37°C and 5% CO2, and the wells were flooded with 3 mL of culture medium. Media was replenished every week for the first two weeks and every 2-3 days after the second week for up to 21 days. Pericytes were also cultured in the presence or absence of 10 µM and 20 µM of Y-27632 (HB2297, Hello Bio Inc, Princeton, NJ), a chemical antagonist of actin-myosin mediated contractility via inhibition of Rho-associated protein kinase (ROCK) dissolved in PBS. Y27632 drug wash out involved pipetting out the existing culture media with the drug, washing the wells twice with 1 mL of the culture media followed by replenishment with 3 mL of fresh culture medium.

### Time-lapse, epi-fluorescence, and confocal microscopy

The 6-well plates were placed in a microscope (AxioObserver Z1, Carl Zeiss, Jena, Germany). Time lapse videos were taken with 20x objective at 5- or 10-min intervals and mosaic images were also taken of the entire scaffold with the same objective. Cells and spheroids were fixed with 4% paraformaldehyde, permeabilized in 0.1% Triton X100 solution and blocked in 5% goat serum. Cells were stained for nuclei using DAPI (Invitrogen), F-actin stress fibers using Phalloidin (Alexa Fluor™ Plus 647 Phalloidin, Invitrogen), while the fibers were labeled prior to seeding via incubation with Rhodamine Fibronectin (Cytoskeleton Inc.) for 1 hour. Labeled spheroids were imaged in a LSM 880 confocal microscope (Carl Zeiss) to obtain fluorescent images and z-stacks.

### RNA isolation and gene expression

Cells and spheroids on fiber networks were lysed at seven-day intervals for three weeks. Total RNA was isolated using NucleoSpin® RNA kit (Takara/Clontech 740955), followed by cDNA synthesis using High Capacity cDNA Reverse Transcription kit (ThermoFisher 4368814) and quantification using Qubit™ ssDNA Assay (ThermoFisher Q10212), all performed according to the manufacturers’ instructions. cDNA inputs were either 10ng (Sm22, Smtn, PPIA) or 100ng (Acta2, Cnn1) for qPCR using Taqman™ technology (ThermoFisher 4369016; refer to Sup. Table 1 for primer/probe details) performed using a QuantStudio™ 3 thermal cycler (ThermoFisher A28131). StepOne Plus v 2.2 software was utilized for data analysis; amount of gene expression normalized to the housekeeping gene PPIA was calculated using the 2^-ΔCt^ method.

### Quantitative analysis of cells and spheroids

Quantitative analysis of the cells and spheroids from the time-lapse videos and images were performed using ImageJ (NIH; https://imagej.nih.gov/ij/). Cell and spheroid boundaries were manually outlined, and area, circularity, and aspect ratios were extracted. Cell and spheroid forces were calculated using our established nanonet force microscopy technology.^27,30,32,33^ This technique involves the use of aligned fiber networks to calculate cellular forces. The thinner aligned fibers fused to the stiff base fibers on both sides are modeled as fixed-fixed beams. The deflection of the fiber by the cell/spheroid is used to calculate the force exerted on the fibers. Quantification of migration rates was performed via centroid tracking. Persistence was calculated as the total displacement of the cell/spheroid divided by the total distance travelled. Polarity of the nuclei within the spheroids were checked against the polarity of the spheroid by manually outlining the nuclear boundary and spheroid boundary from confocal z-stack images. Differences between the major axes of both boundaries were calculated as an angle between 0° and 90°. Student’s t-tests were used for statistical analysis, with a p value < 0.05 classified as *, <0.01 as **, and <0.001 as ***.

## Results

### Pericytes aggregate into spheroids on suspended fibers

On suspended fiber networks, we found that pericytes self-assembled into multi-cellular aggregates (e.g., spheroids) over timescale of hours (**Figure 1(a)**, Sup. Movie 1-3). We designed the fiber networks such that individual fibers maintained their integrity (did not merge with other fibers) by depositing a parallel array of small diameter fibers (200, 500, and 800 nm) on top of large diameter generally non-deformable fibers (∼2000 nm termed *base* fibers). The two layers of the fibers were fused at the contact points resulting in fixed-fixed boundary conditions.^28,30^ Cells applied contractile forces as visualized by the deflection of the fibers, resulting in the assembly of spheroids. To visualize spheroid 3D organization on fiber networks using confocal microscopy, pericytes were seeded on fibers coated with Rhodamine-conjugated fibronectin. Confocal microscopy of DAPI and phalloidin-labeled cells revealed that spheroids formed around multiple fibers (above and below the plane of fibers, **Figure 1(b)**). Spheroids were classified as having large areas (∼6187.63 ± 3449.76 μm^2^) and high circularities (∼0.71±0.14) (**Figure 1(c)**) and were significantly larger (P<.001, Sup. Figure 1(b,c)) from single cells or small clusters of three or fewer cells, which have smaller areas (1066.35 ± 832.35 μm^2^) and lower circularities (0.21 ± 0.08). We found that as the area of the spheroids increased, the number of cells within them increased linearly (**Figure 1(d)**). The nuclei of cells within spheroids were observed to have a polarity which matched the polarity of the spheroid, with the angle between the major axes of the ∼50% of nuclei and the spheroid being within 30° (**Figure 1(e)**).

**Figure 1:**
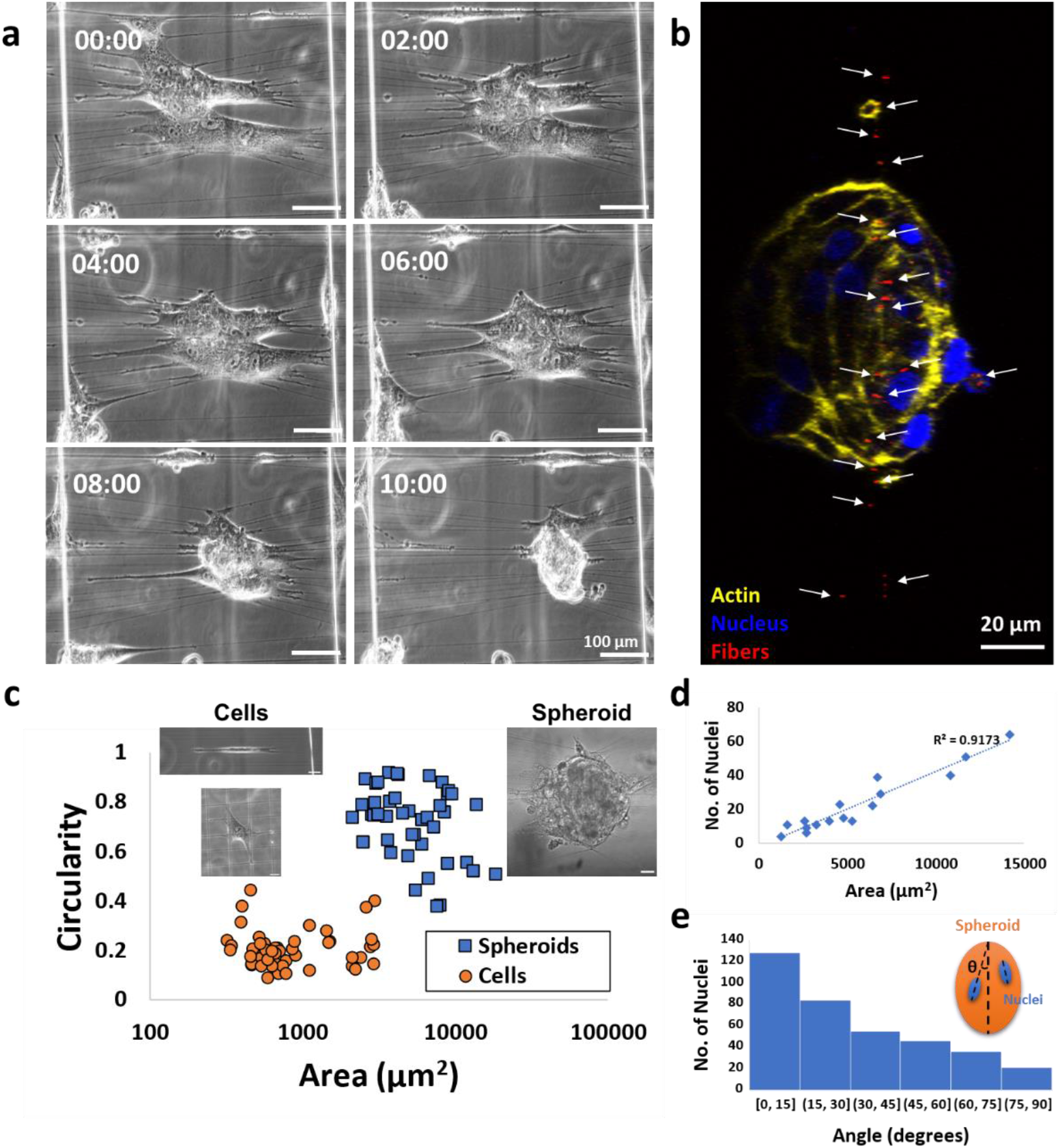
Spheroid architecture on fiber networks: (a) Time lapse images of spheroid assembly from a group of individual pericytes (Scale bar: 100 µm) (b) Confocal image of a side view of a spheroid. The fibers (in red, indicated with white arrows) are present within the spheroids. (Red - Fibers; Yellow – Actin; Blue – Nucleus; Scale bar: 20 µm). (c) Cells on fiber networks (inset, left) had lower circularities and smaller areas while spheroids (inset, right) were classified with high circularities and larger areas. (N=49, Inset scale bars: 50 µm) (d) The number of nuclei as a marker for cell number increased linearly with increase in spheroid area. (N=19) (e) Nuclei within the spheroids have their major axes aligned to the major axis of the spheroid, indicated by the large number of nuclei having a small difference between the angles of the major axes of the spheroid and the nucleus (N=391 (nuclei), N=19 (spheroids))

### Biophysical cues from the underlying microenvironment influence spheroid assembly

We next inquired if spheroid assembly on fibers was regulated by the size and architecture of fiber networks. To answer this, we tracked pericytes cultured on STEP fiber networks of three diameters and two architectures for three weeks. We found that spheroids formed within 3-7 days of cell seeding on aligned fiber networks for all three fiber diameters (**Figure 2(b)(i)-(iii)**, Sup. Figures 2-4). The number of spheroids and their size notably increased during the second week of culture. In multiple instances, we found that cells and cell clusters homed to the large ∼2000 nm *base* fibers and merged to form elongated structures around and along *base* fibers. These cellular aggregates were found to be of larger areas (158,806±42,875 μm^2^) and lower circularities (0.37±0.02) (**Figure 3(a)**, Sup. Figure 2-4). On 200 nm aligned fiber networks, some elongated structures contracted along their major axis into large, individual spheroids during the third week concurrent with fiber remodeling; parallel fiber fused-junctions with *the base* fiber were observed to break due to increased contractility resulting in merging of parallel fibers (Sup. Movie 4-6, Sup. Figure 9). Interestingly, on 500 nm aligned fiber networks, we observed elongated structures to be of larger areas (2,295,227±1,673,914 μm^2^) and low circularities (0.34±0.18) that persisted from the second week of culture (**Figure *2*(b)(ii)**, Sup Figure 3, **Figure 3(a)**). In the third week of culture unlike the 200 nm and 800 nm fiber diameter counterparts, these structures did not change in size or shape, though new spheroids were identified (Sup Figure 3).

**Figure 2:**
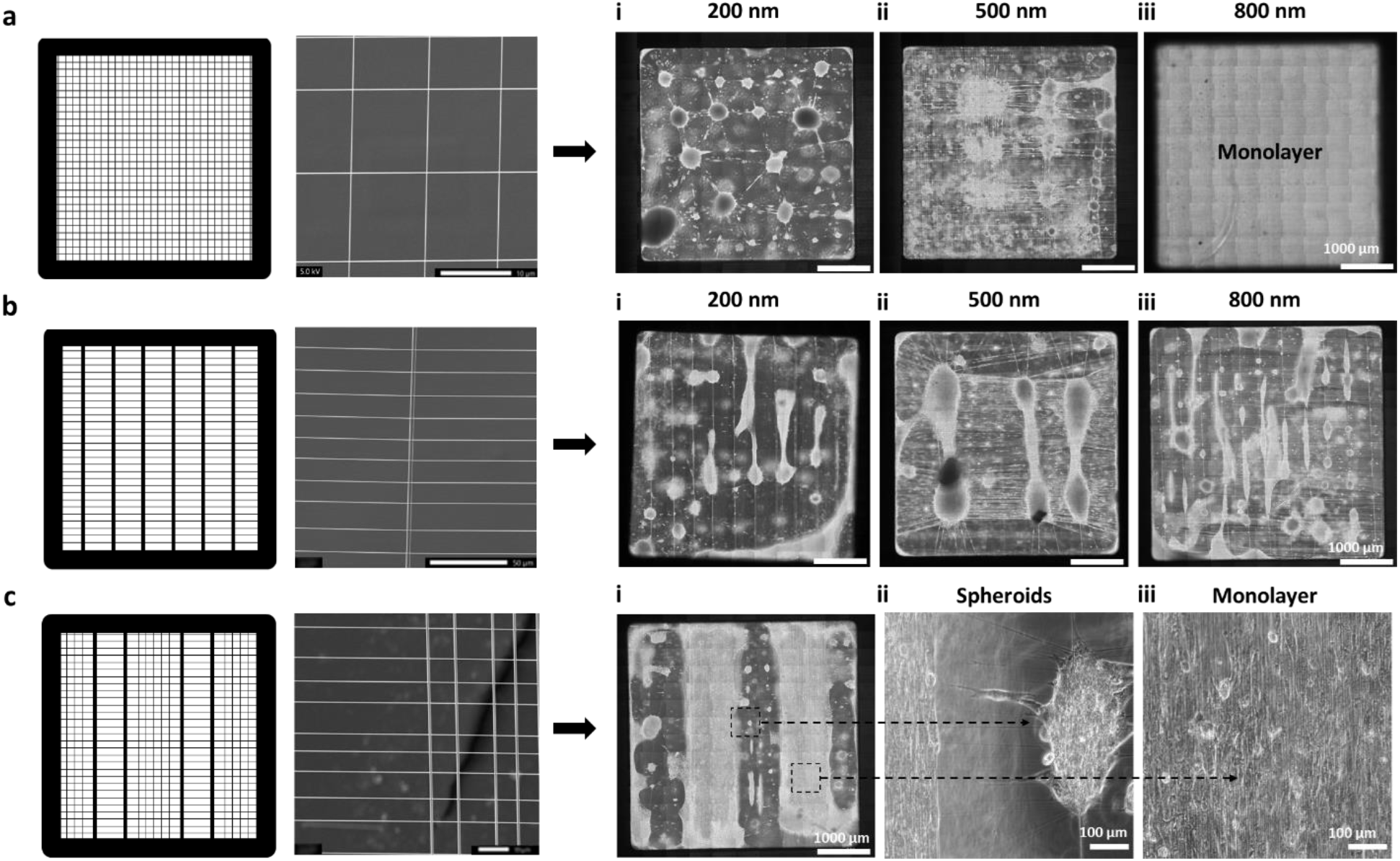
Fiber diameter and architecture influence pericyte spheroids: Pericyte cultures grown on (a) crosshatch fiber scaffolds and (b) aligned fiber scaffolds with (i) 200, (ii) 500 and (iii) 800 nm fiber diameters after two weeks in culture. Crosshatch fibers show a strong influence of fiber diameter for spheroid assembly while aligned fibers are not affected by different fiber diameters. (c) (i) Pericyte cultures grown on fiber scaffolds with both crosshatch and aligned regions on the same scaffold after two weeks in culture. Regions with aligned fiber architectures show spheroid formations (ii) while those on crosshatch fiber networks grow in monolayers (iii).

**Figure 3:**
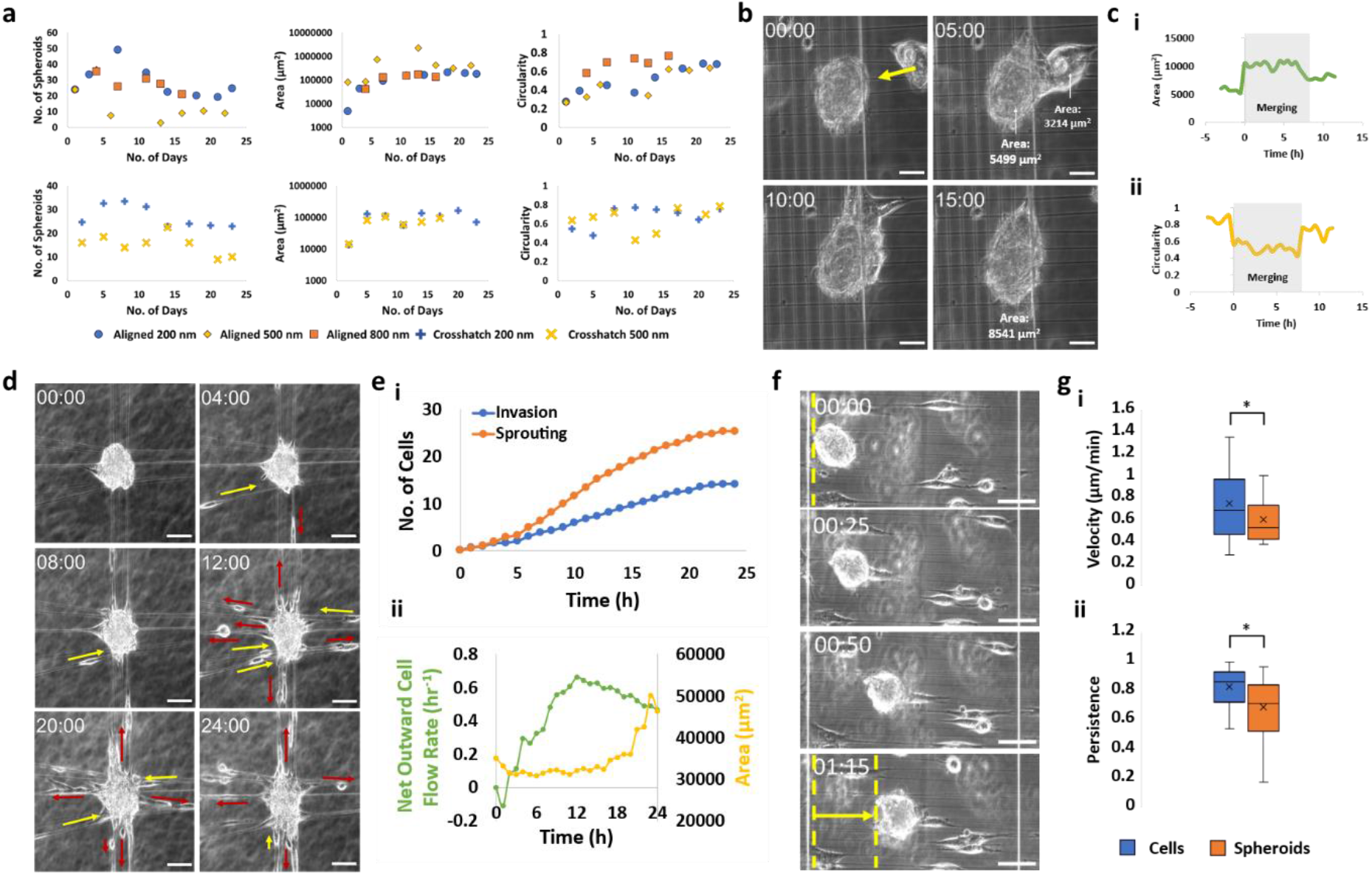
Spheroid interactions with fiber networks: (a) Changes in the number of spheroids, area, and circularity of spheroids with time when cultured on suspended fibers of varying diameter (200, 500 & 800 nm) and architecture (force and crosshatch fiber networks). (b) Two spheroids merging. (Scale Bar: 50 μm) (c) Changes in area (i) and circularity (ii) of the large (left) spheroid in (b) during merging. (d) Time lapse images of cells continuously invading into (yellow arrows) and sprouting out of (red arrows) spheroids. (Scale Bar: 100 μm) (e) (i) Cells sprouting out of spheroids occurred with a higher frequency than cells invading into the spheroid (N=18). (ii) Changes in outward cell flow rate and area with time in spheroids with invading and sprouting cells (N=18). (f) Time lapse images of spheroids moving along the aligned fibers. (Scale Bar: 100 μm) (g) Average velocity (i) and persistence (ii) of migrating spheroids. (N=23)

Next, we assessed spheroid assembly on different fiber architectures (e.g., crosshatch network of fibers. We designed the crosshatch network of fibers by depositing two layers of same diameter fibers orthogonal to each other and fused at the intersections. We found a strong influence on fiber diameter in the crosshatch fiber configuration on spheroid assembly. Among the three diameters, 200 nm crosshatches were the easiest for pericytes to form spheroids (Figure *2*(a)(i)). The spheroids deformed the fiber networks during culture, breaking the fused fiber junctions and causing the fibers to align in a direction perpendicular to their boundary (Figure *2*(a)(i), Sup. Figure 9). Multiple spheroids were observed across the entire scaffold, with multiple fiber interconnections connecting them (Sup. Figures 5-6). These spheroids exhibited high circularity values (0.65 ± 0.02) throughout their three weeks in culture (Sup Figure 5-6), unlike aligned fiber networks, where long structures were predominant at the base fibers during the second week which then contracted into spheroids. Fewer spheroid occurrences were observed on 500 nm crosshatch fiber networks. (Figure *2*(a)(ii), Sup. Figure 6). Regions where pericytes were observed to remodel fibers were associated with higher occurrences of spheroid assembly when compared with regions exhibiting intact fibers, where cells grew into monolayers. Interestingly, pericytes cultured on 800 nm crosshatch networks did not form any spheroids (Figure *2*(a)(iii)). Monolayers were formed over long-term cultures, with the fiber networks maintaining the crosshatched architecture (Figure *2*(a)(iii), Sup. Figure 7).

Designing fiber scaffolds of precise diameters and architectures provided us a unique opportunity to consider control of the spatial patterning of spheroids. To this end, we reasoned that a design of 800 nm crosshatches interspaced with parallel 200 nm diameter fibers would result in monolayers and spheroids, respectively. Indeed, we observed spheroids formed only in the aligned fiber regions, and monolayers formed in regions having crosshatched fiber networks (**Figure *2*(c)(i-iii)**, Sup. Figure 8), thus providing new abilities to pattern spheroid assembly using suspended fiber networks.

Next, from time lapse images captured every 2-3 days, we calculated the dynamics of spheroid assembly. Overall, design of fiber networks yielded new knowledge in the ability of cells to aggregate into spheroids and spatially pattern them.

### Spheroids exhibit dynamic interactions with their surrounding microenvironment

Next, we observed how spheroids dynamically interacted with other cells and their microenvironment using timelapse imaging over short intervals (∼8-24 hours). On aligned fibers networks, we observed an unusual sliding/movement of spheroids along the fibers (**Figure 3(f)**, Sup. Movie 7-9) with an average velocity of 0.60 ± 0.22 μm/min (**Figure 3(g)(i)**) and an average persistence (the ability of the cell/spheroid to maintain the direct of motion) of 0.69 ± 0.2 (N=23) (**Figure 3(g)(ii)**). Spheroids on the aligned fiber networks tended to migrate from the 200 nm aligned regions towards the larger 2 μm *base* fibers (Sup Movie 7,8) and used these stiffer fibers as a hub to grow in size, as they remained predominantly stationary on the *base* fibers (Sup. Movie 10-12). We also observed individual cells sprouting from and migrating away from the spheroids or migrating towards and joining these spheroids ((**Figure 3(d)**, Sup. Movie 13-15). The number of cells leaving the spheroid was found to be larger than the number of cells joining the spheroid (N=18) (**Figure 3(e)(i)**). Over a period of 12 hours, we calculated the net outward cell movement, which is the cumulative difference between the cells sprouting away with those joining the spheroid. We found that the net outward cell movement rapidly increased, while the spheroid area remained constant. After 12 hours, we found the cell movement rate decreased gradually which was concomitant with increase in area increase (N=18) (**Figure 3(e)(ii)**). On both aligned and crosshatched fiber networks, spheroids were found to remodel the fiber arrangement during their assembly and growth. Interestingly, in certain instances, we observed exchange of cells in spheroid-spheroid interactions between spheroids in different locations via connecting fibers between them (Sup. Movie 4-6, Sup. Figure 9). In other remarkable instances, we observed spheroid mergers (**Figure 3(b,c)**, Sup. Movie 16-18) that contributed to their increase in size. On aligned fiber networks, multiple cells and spheroids accumulated in different regions along the base fibers to form elongated structures (Sup. Figure 2-4). Larger spheroids and elongated structures exhibited were together capable of exerting large forces to deform *base* fibers and exhibit multiple interactions with other cells or fibers (Sup. Movie 4-6). Overall, we quantitated multiple cell-spheroid, spheroid-spheroid, and spheroid-fiber interactions that either led to increase in size or change in shape of spheroids.

### Forces and contractility in spheroids

Since pericyte spheroid assembly caused the fiber networks to deflect, we measured the forces exerted during this process. Using nanonet force microscopy^27,30,32,33^, we found the spheroid force exertion to be higher (P=.003) than individual pericyte forces. Individual cells had an average force of 41.55 ± 18.78 nN (N=10), while spheroids had a wide range of forces depending on their size (1383 ± 1787 nN ((N=49), Sup. Figure 10). Spheroid forces increased with area (**Figure 4(a(i))**). A trend of decreasing force to 962 ± 1396.69 nN from 1882 ± 2098.17 nN when individual cells sprouted from the spheroid (P=.09), while area reduced to 6428 ± 1985.40 µm^2^ from 7343 ± 2732.68 µm^2^ (P=.02, N=12, **Figure 4(a)(ii,iii)**). To study the forces involved during the assembly of spheroids, transient forces were calculated from the point when multiple individual cells came into contact with each other until they assembled into spheroids. As the collective area occupied by the cells decreased, and the cells became rounded and migrated closer to one another (Sup. Movie 1-3), the average forces reduced from 1766 nN during the start of assembly to 327 nN when spheroids were fully formed while area reduced by 47% from 16529 μm^2^ to 8654 μm^2^ (N=10, **Figure 4(a)(iv)**).

**Figure 4:**
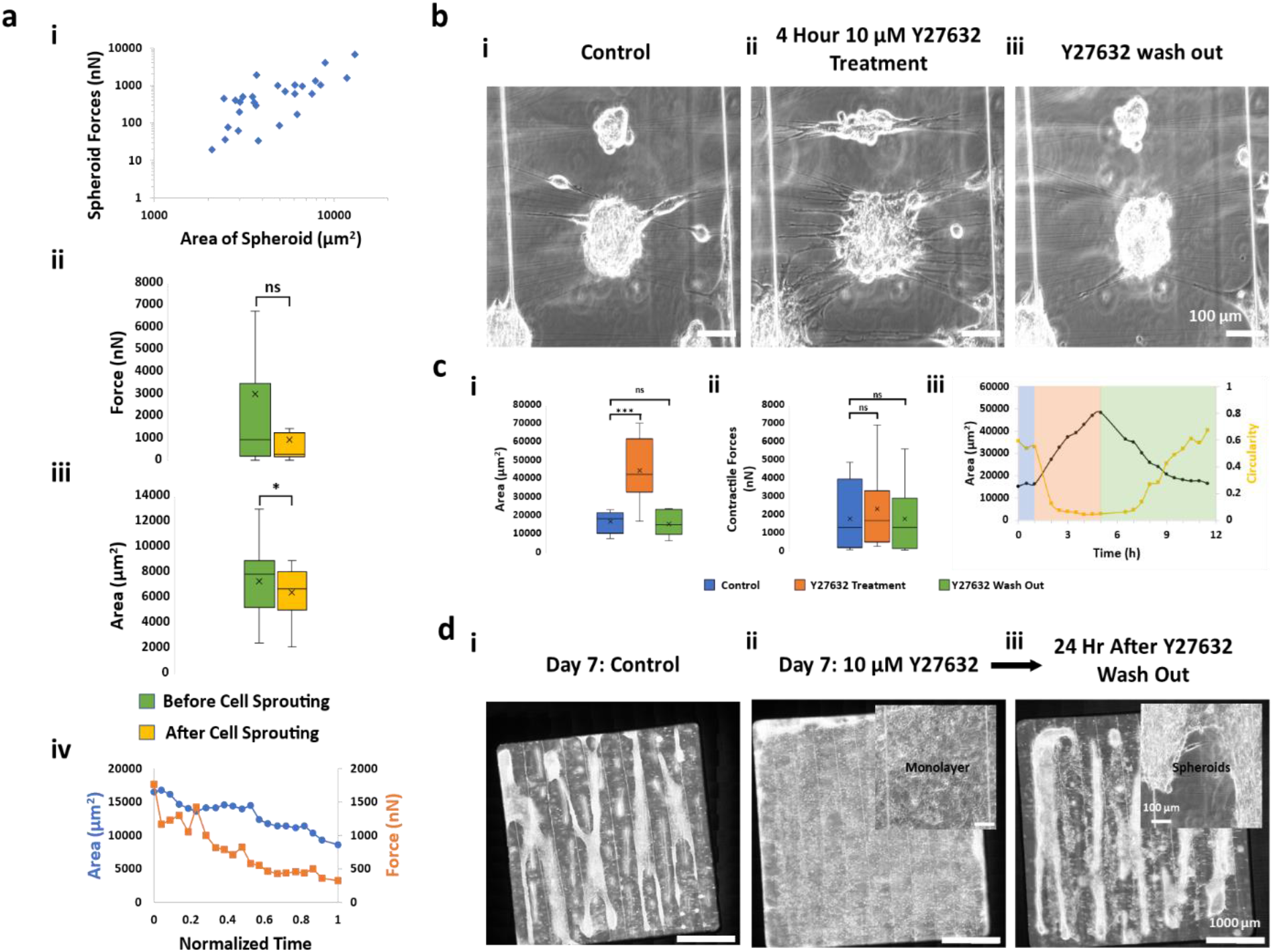
Forces and contractility play an important role in spheroids: (a) (i) Relationship of spheroid forces with spheroid area (N=27). (ii) Contractile forces and (iii) spheroid area before and after an individual cell sprouts from the spheroid (N=12). (iv) Dynamic changes in contractile forces and area during spheroid assembly (N=9). (b) (i) Spheroids before drug treatment (ii) Effect of treatment of 10 µM Y27632 (ROCK-inhibitor) for 4 hours on spheroids and (iii) recovery of the spheroids after drug wash off (Scale bars: 100 µm). (c) Changes in (i) area and (ii) contractile forces during Y27632 treatment and after Y27632 wash out (N=10). (iii) Dynamic changes in area and circularity of the spheroids during Y27632 treatment and after Y27632 wash out (N=10) (d) Images of the suspended fiber networks after seven days of (i) control cells and (ii) cells seeded with Y27632 (inset shows a magnified region of the scaffold). (iii) Spheroid/organoid formation seen after 24 hours of Y27632 wash out. Insets shows a magnified area of the fiber network. (Scale bars: 1000 µm (inset: 100 µm))

**Figure 5:**
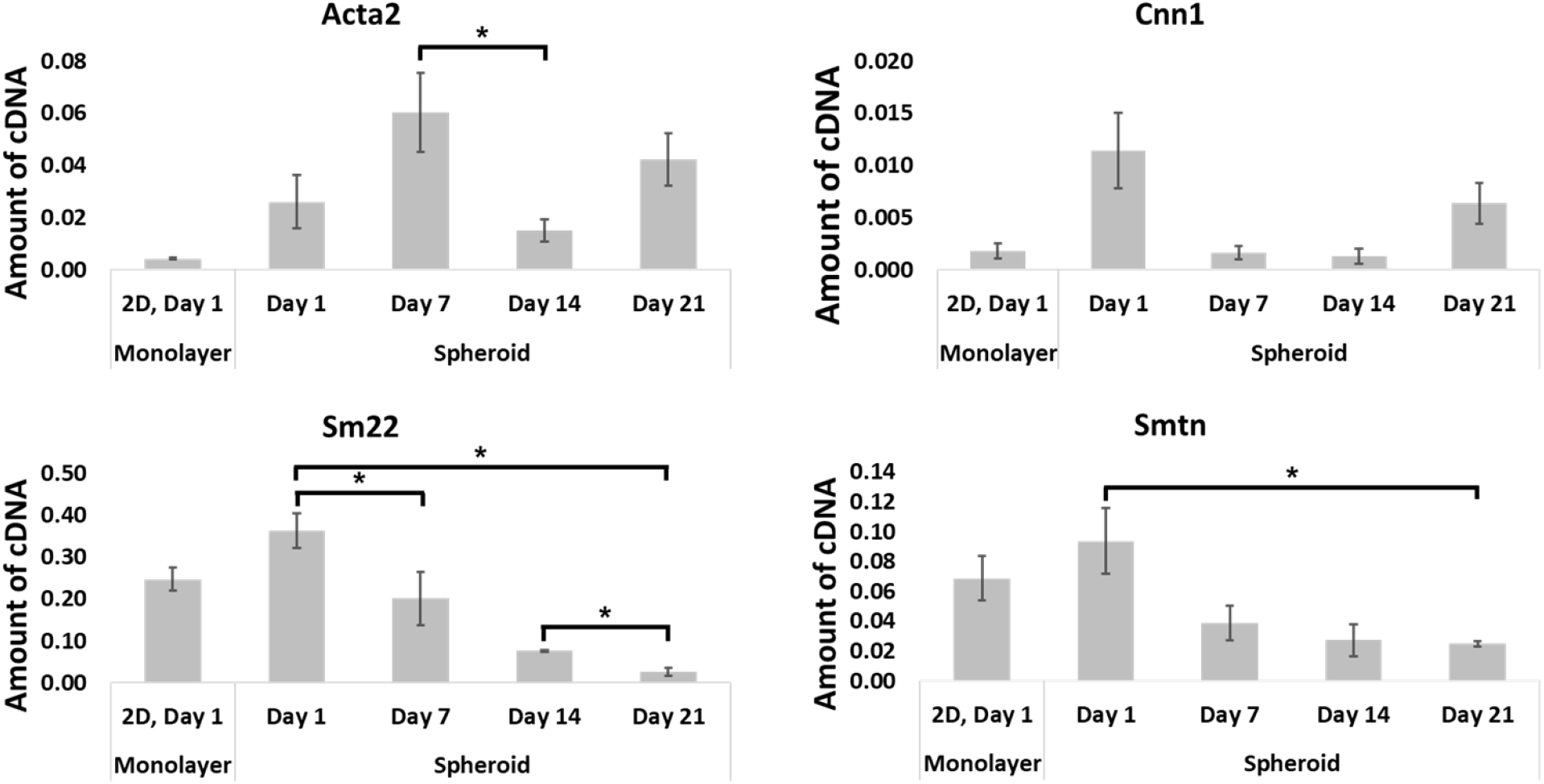
Acta2, Cnn1, Sm22, and Smtn gene expressions in pericytes plated on either 2D plates (collected on Day 1) or on fiber networks (collected on Day 1, Day 7, Day 14, and Day 21 post-seeding).

Next, we interrogated the role of cell contractility in spheroid assembly by pharmacologically inhibiting ROCK using Y27632. We observed that spheroids rapidly (within 1-2 hrs) lost their compactness and migrated away from the spheroid along the aligned fibers without apparent loss of cell-cell adhesion (**Figure 4 (b)(i,ii)**), Sup. Movie 19-21) evidenced by an increase in the area and a decrease in the circularity (**Figure 4(c)(i,iii)**). However, addition of Y27632 did not affect the spheroid force exertion on fibers (N=10) (Figure 4(c(ii))). The impact of ROCK inhibition was reversible. Restoring control conditions by washing the cells and replenishing with fresh culture medium after 4 hours caused the cells to quickly reform spheroids after 6-8 hours (**Figure 4 (b)(iii)** and **Figure 4(c)(i-iii)**). Increasing the dose from 10 μM to 20 µM and treatment time from 4 hours to 8 and 15 hours produced similar effects with cells moving further away from the spheroids with time while preserving cell-cell adhesions (Sup. Movie 22-26). ROCK inhibition at the time of cell seeding prevented spheroid assembly caused the pericytes to grow in monolayers, while control cultures lacking Y27632 showed multiple instances of spheroid assembly and elongated structures (**Figure 4(d)(i-ii)**, Sup. Figure 11). Reversal by drug wash off showed cells immediately contracting towards each other at multiple locations to form spheroids and elongated structures within 24h (**Figure 4(d)(iii)**), Sup. Movie 27-29), as seen in the control culture.

Next, we measured gene expression to better understand the dynamics of spheroid assembly. We compared contractility markers *Acta2, Cnn1, Sm22* and *Smtn* from RNA isolated from spheroids on fiber networks on days 1, 7, 14, and 21. None of the genes examined showed a significant change at day-1 post-seeding on fiber networks compared with 2D monolayer culture (P=0.059). However, while *Acta2* expression appeared to increase over 7 days, it also displayed a biphasic expression pattern, and decreased between days-7 and days-14 on fiber networks (0.060 ± 0.015 vs 0.015 ± 0.004, P=0.045). *Cnn1* expression showed a trend of downregulation approaching statistical significance noted at both day 7 and day 14 when compared with day 1 (0.011 ± 0.004 vs 0.002 ± 0.001, P=0.056 and 0.011 ± 0.004 vs 0.001 ± 0.001, P=0.051, respectively); however, these differences did not reach statistical significance within 5% confidence. *Sm22* expression decreased with time on fiber networks when compared with the first day of seeding on the scaffolds (0.076 ± 0.002 vs 0.363 ± 0.042, P=0.002 and 0.026 ± 0.010 vs 0.363 ± 0.042, P=0.001, respectively). Additionally, day 21 spheroids exhibited decreased *Sm22* when compared with day 14 (0.026 ± 0.010 vs 0.076 ± 0.002, P=0.008). By 21 days post seeding, *Smtn* gene expression decreased when compared with day 1 gene expression (0.025 ± 0.002 vs 0.094 ± 0.022, P=0.036). Additionally, day 7 and day 14 showed a trend for decreasing *Smtn* gene expression when compared with day 1 (0.039 ± 0.012 vs 0.094 ± 0.022, P=0.092 and 0.027 ± 0.011 vs 0.094 ± 0.022, P=0.054), which did not reach statistical significance within 5% confidence. Overall, we report modulation of force exertion during spheroid assembly and a role for cell contractility in spheroid dynamics over long term culture, reflected through changes in expression of genes involved in contractility.

## Discussions

Mechanical cues from the underlying microenvironment influence various aspects of cellular behavior.^34,35^ The native ECM of tissues is comprised of protein fibers with varying diameter and stiffness^36,37^ which are extensively remodeled for various processes such as tissue homeostasis,^38^ cell migration,^39^ and tumor progression.^40–42^ Matrix stiffness has been shown to affect spheroid morphology and behavior such as disaggregation in ovarian cancer cell spheroids^43^ and increased invasiveness of pericytes and melanoma cell spheroids.^20,23^ Here, we demonstrate that biophysical cues such as changing the fiber diameter and fiber architecture play a key role in dynamically influencing assembly, growth, and behavior of pericyte spheroids.

It is well appreciated that the mechanical properties of spheroids can be influenced by fibrous matrices. For example, malignant breast cancer cell spheroids, when treated with collagenase to degrade collagen showed an increase in area and decrease in stiffness.^44^ Similarly, previous studies with hepatoma cell spheroid assembly have demonstrated the role of ECM and integrin-ECM binding in promoting spheroid assembly for rapid aggregation followed by E-cadherin upregulation and cell-cell adhesion.^22,45^ In the present study, we evaluated spheroid assembly by pericytes and found that fiber diameter and architecture promoted spheroid assembly through fiber remodeling. In aligned fibers, changes in the diameter influenced the way in which cells aggregated together, leading to aggregates of different sizes and morphologies (spheroids and elongated structures). Unexpectedly, large diameter crosshatch networks did not support the assembly of spheroids, presumably due to the high stiffness of the fiber scaffolds which prevented fiber remodeling. Thus, in scaffolds incorporating smaller diameter parallel (e.g., aligned) arrangements of fibers juxtaposed with large diameter crosshatches yielded zones of spheroids and monolayers, respectively. Overall, extensive fiber remodeling was observed around spheroid occurrences on fiber networks, while regions/scaffolds without spheroids maintained an intact non-deformed fiber architecture. Thus, the fiber architecture, stiffness (fiber diameter), and the cells’ ability to remodel the fibers seem to be critically linked to spheroid assembly.

Various processes such as morphogenesis, wound healing, and tumors involve a mode of cell migration known as collective cell migration.^46–49^ Beaune *et al*.^50^ characterized different forms of collective cell migration with murine sarcoma cells on rigid substrates based on their morphology during migration. Previously, we have shown that leader and follower cells in single, chain, and collective modes invade from a monolayer interfaced with parallel and crosshatch network of fibers^51^. Here, on aligned fiber networks, we found a remarkable ability of some spheroids being dragged or pulled by leader cells during migration. This form of migration by one or more leader cells is similar to collective cell migration initiated by leader/tip cells.^52–55^ We also saw some instances of spheroids moving without a leader cell, possibly due to imaging challenges in locating the leader cell or a yet unclear mode of migration utilizing some form of coordinated internal signaling within spheroids to move as a whole. A puzzling component in spheroid migration is how were the spheroids able to migrate? Confocal images show that the spheroids were formed on either side of fibers (fibers were inside the spheroids). Leader cells being able to pull the spheroids suggests a sliding mode that would only be possible on parallel arrangement of fibers similar to beads on a string. Indeed, spheroids were found to be non-migratory on the larger diameter (stiffer) orthogonal *base* fibers on aligned fiber networks, and on crosshatch network of fibers. While the biological origins of spheroid migration are not clear, our observations clearly indicate an influence of both stiffness and fiber architecture on spheroid migration.

The mechanical force exertions by spheroids and responses when external forces are applied on it are a focal point of active research.^21,56–58^ We calculated the dynamic forces exerted by the spheroid on the fiber scaffolds during various behaviors. During initial stages of spheroid assembly, individual cells co-exist in the same 2D plane, and collectively exert a larger force on the scaffold. With spheroid assembly, a continuous and considerable drop in force exertion was observed as some cells detach from the fibers and assemble into 3D structures. As the cells contract into spheroids, they tend to pull the fibers with them. Stiffness, matrix alignment, and tension in the fibers have been shown to increase invasiveness and sprouting in endothelial and melanoma cell spheroids.^19,23,59^ Our fiber networks include highly aligned fibers in one or more directions, with the fibers in tension, and represent an effective tool to study individual cells merging into and sprouting out of spheroids. Here, we found a larger number of cells sprouting away from the spheroid when compared to cells merging into the spheroid. Initially, the area of the spheroid was constant even with a large outward flow suggesting that the cells within the spheroid may be dividing at a similar rate. We also show a decrease in spheroid forces when single cells sprout out of the spheroid, indicating the role of individual cells to collectively contribute to spheroid forces.

Actin-myosin contractility can alter spheroid behavior and compaction in various cell types such as human mesenchymal stem cells, breast and ovarian cancer cell lines.^43,60,61^ We confirm a destabilizing effect on pericyte spheroids with ROCK inhibition and show that assembly of spheroids was prevented by inhibiting actin-myosin contractility. Interestingly, while control cells form the spheroids and elongated structures with 7 days, pericytes cultured for a week under ROCK-inhibition failed to assemble into spheroids but immediately did so within 24 hours upon removal of the ROCK inhibitor. Our data suggests a critical role of ROCK-mediated actomyosin contractility in spheroid assembly. The biphasic pattern in expression of genes involved with pericyte contractile function over long term culture of pericyte spheroids on nanofibers suggests that response to biophysical cues are dynamic and further influenced by changes in spheroid size and structure over time. Because pericytes wrap around blood vessels in the microvasculature and their contractility helps restructure vessel wall and regulate fluid flow,^13^ their contractile function is likely influenced by matrix biophysical cues. Here, we show that contractility plays a major role in pericytes’ ability to form and assemble into 3D structures and with inhibition, these behaviors are blocked. Thus, fiber matrices differentially influence pericyte behavior which could have implications for physiological and pathophysiological angiogenesis.

Overall, we demonstrated that aligned and suspended fiber networks in various configurations of differing architecture and diameter (stiffness) enable determination of how biophysical cues can influence pericyte behavior to form either 3D spheroids or 2D cell monolayers. Pericytes sense fiber diameter, architecture, and cause fiber remodeling to adopt a morphology and cues from their underlying microenvironment influence their growth. Our ongoing work is focused on the dynamic interactions of pericyte spheroids on suspended fiber networks to decipher the underlying biology of how stiffness and architecture dictate spheroid behaviors, longevity, and putative translational applications.

## Acknowledgements

SS and ASN thank members of the Spinneret-based Tunable Engineered Parameters (STEP) Lab, Virginia Tech for helpful suggestions and discussions. SS and ASN thank the Institute of Critical Technologies and Sciences (ICTAS) and Macromolecules Innovation Institute at Virginia Tech for the support to conduct this study. The authors recognize and thank Kristin Valchar, Dr. Ibrahim Sultan, and Dr. David Kaczorowski for their assistance with patient enrollment and aortic tissue procurement. ASN acknowledges partial support for this study from the National Science Foundation (NSF grant 1762634, 2119949 and 2107332). JAP acknowledges partial support for this study from the NHLBI award #R01HL131632, the UPMC Pellegrini Chair in Cardiothoracic Surgery, University of Pittsburgh Physicians, and the Department of Cardiothoracic Surgery, University of Pittsburgh School of Medicine.

## Author contributions

ASN and JAP designed and supervised the project. SS fabricated fiber scaffolds, conducted experiments and analyzed data with help from ASN and JAP. JAP and JCH isolated cells and performed gene expression studies. SS wrote the initial draft of the manuscript. All authors edited and finalized the manuscript.

## SUPPLEMENTARY FIGURES

**Supplementary Figure 1:**
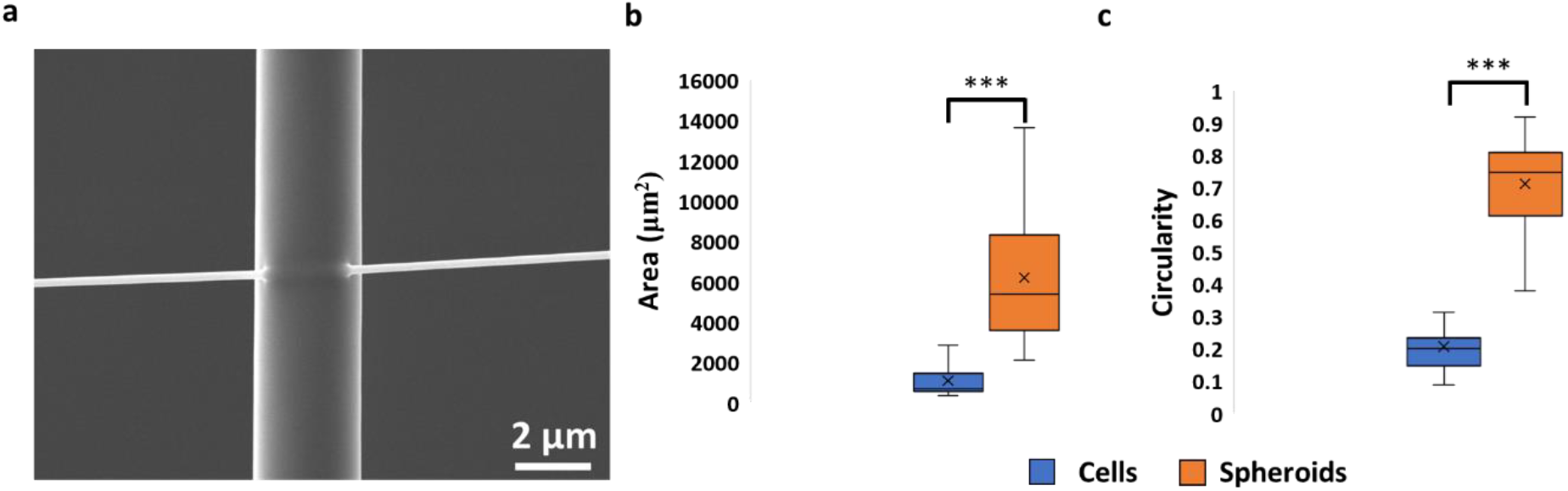
(a) Fused junctions at the intersections of the fiber networks. Difference in (b) area and (c) circularity between cells and spheroids (N=49)

**Supplementary Figure 2:**
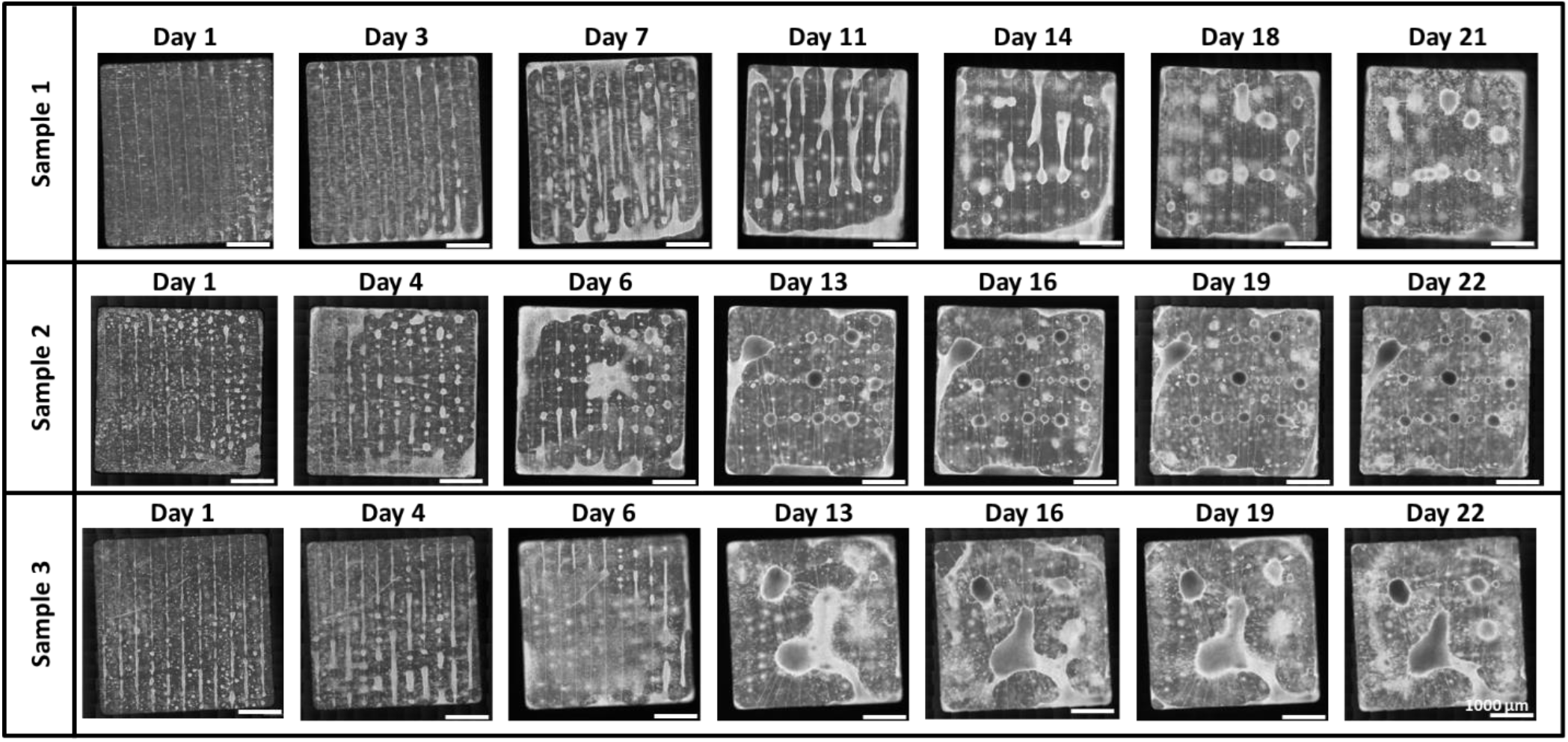
Mosaic images of pericyte cultures on aligned fiber networks with 200 nm aligned fibers and 2 µm base fibers cultured for ∼3 weeks (Scale bars: 1000 µm).

**Supplementary Figure 3:**
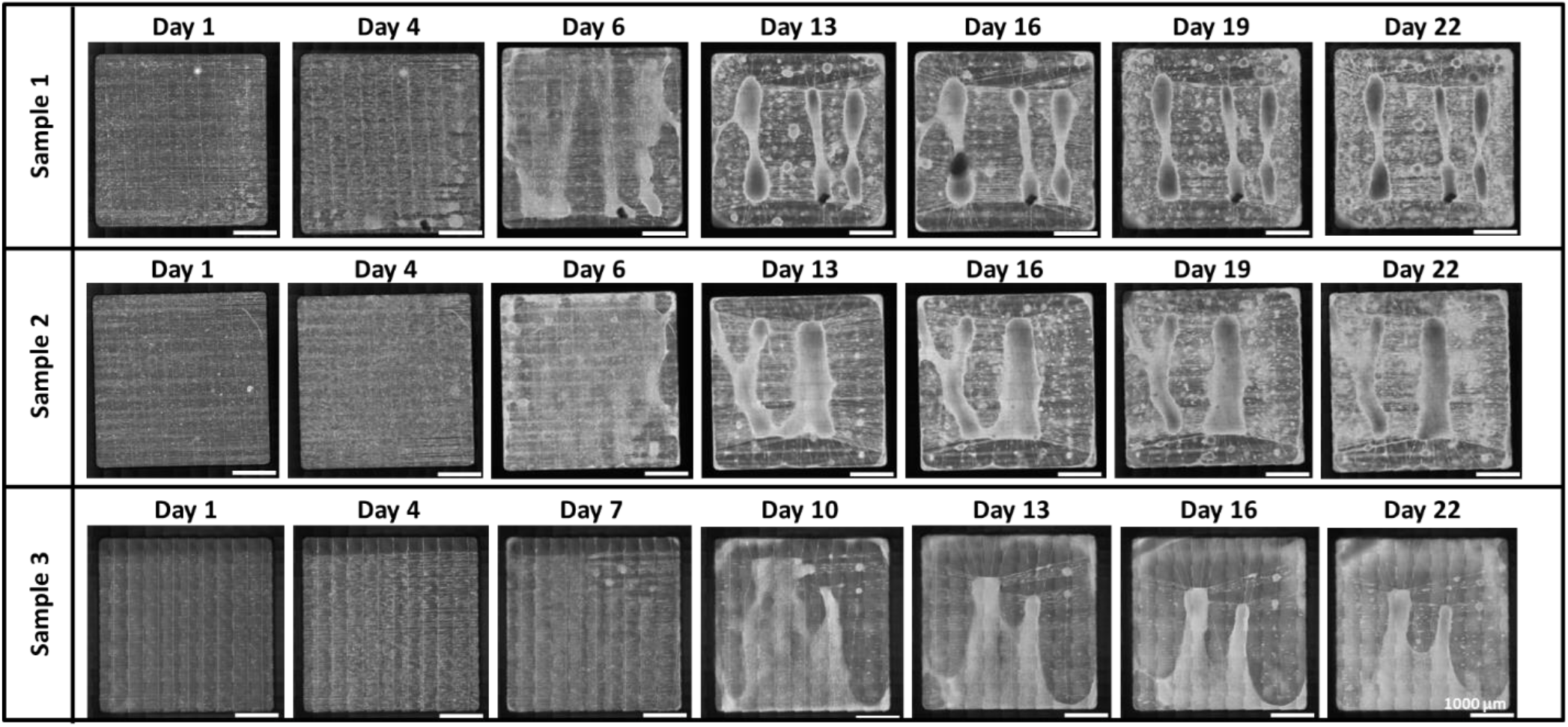
Mosaic images of pericyte cultures on aligned fiber networks with 500 nm aligned fibers and 2 µm base fibers cultured for ∼3 weeks (Scale bars: 1000 µm).

**Supplementary Figure 4:**
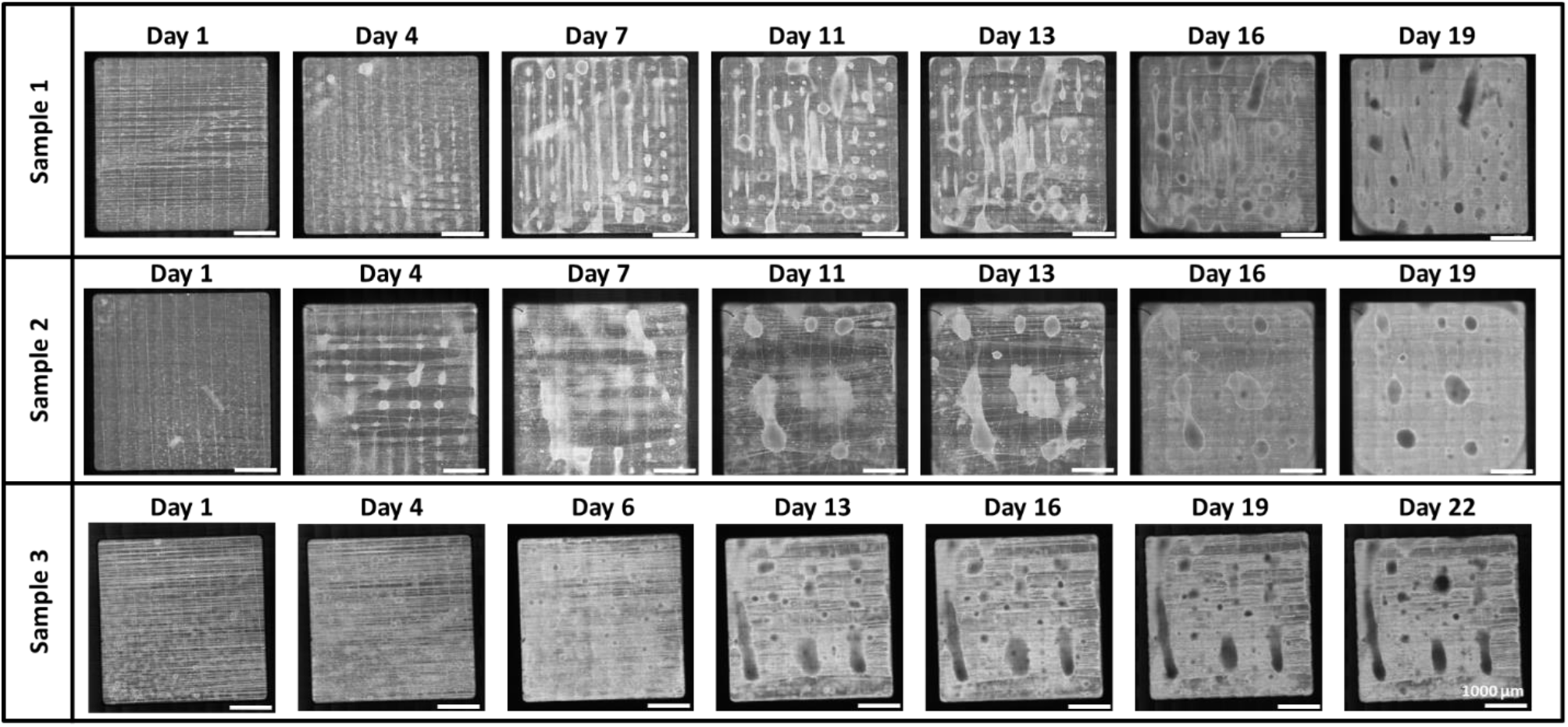
Mosaic images of pericyte cultures on aligned fiber networks with 800 nm aligned fibers and 2 µm base fibers cultured for ∼3 weeks (Scale bars: 1000 µm).

**Supplementary Figure 5:**
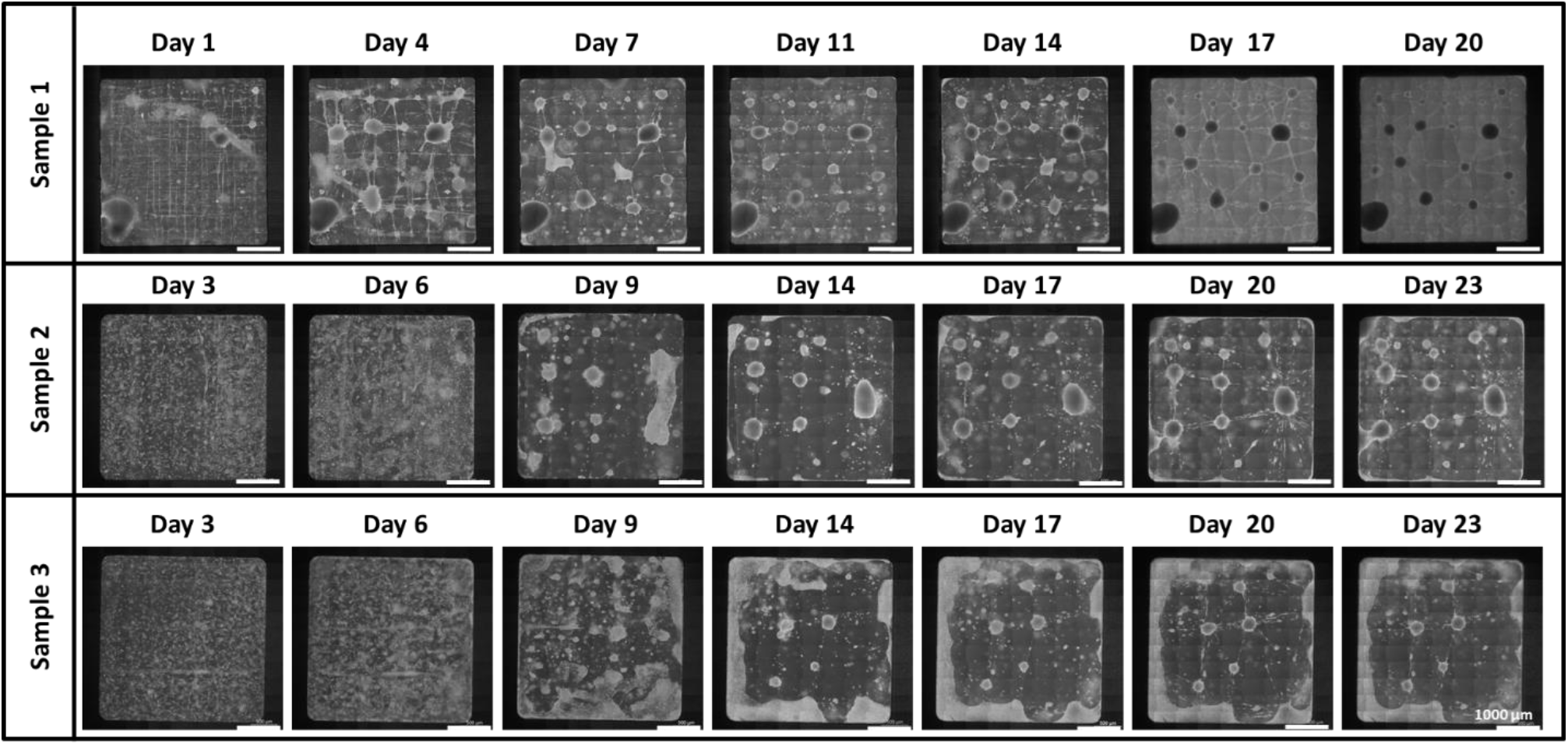
Mosaic images of pericyte cultures on crosshatch fiber networks with 200 nm fibers cultured for ∼3 weeks. (Scale bars: 1000 µm).

**Supplementary Figure 6:**
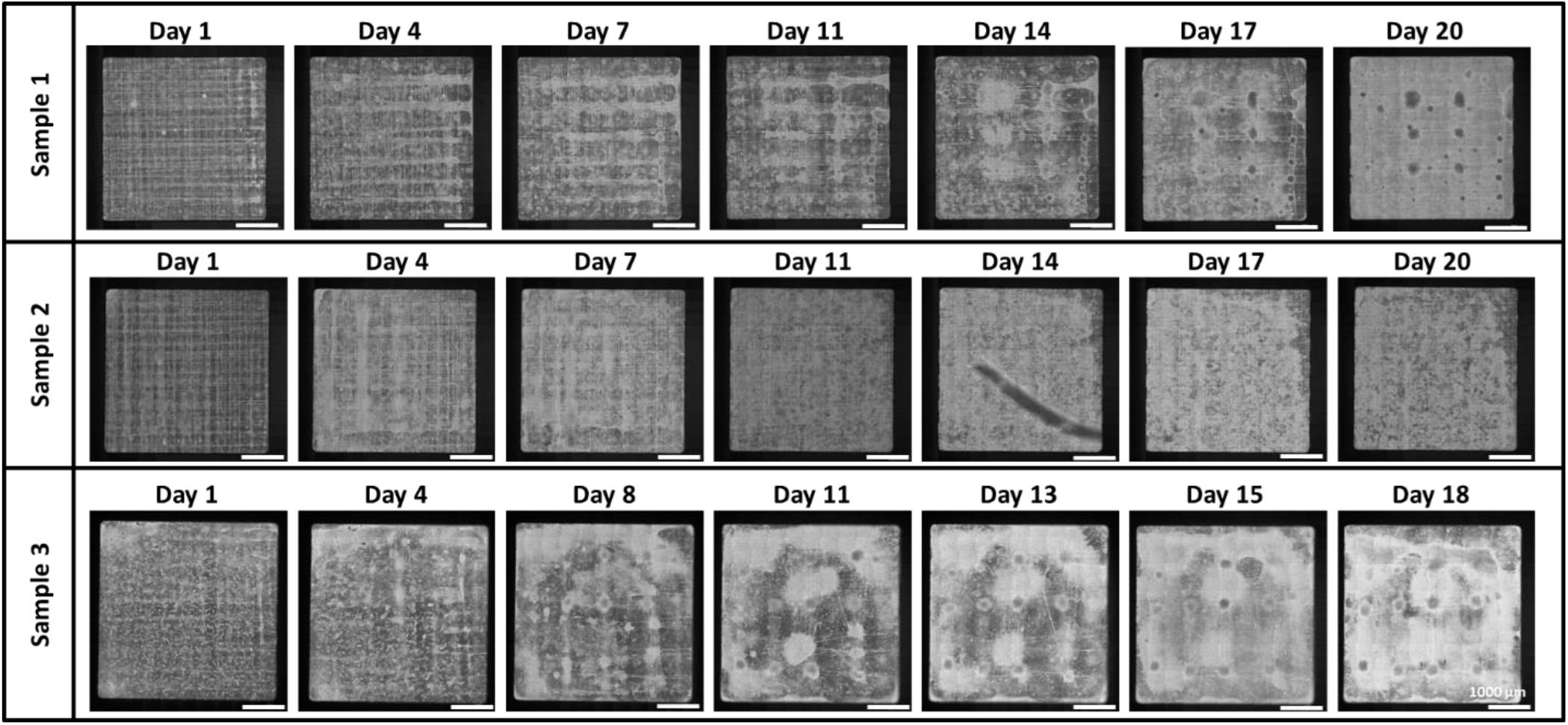
Mosaic images of pericyte cultures on crosshatch fiber networks with 500 nm fibers cultured for ∼3 weeks. (Scale bars: 1000 µm).

**Supplementary Figure 7:**
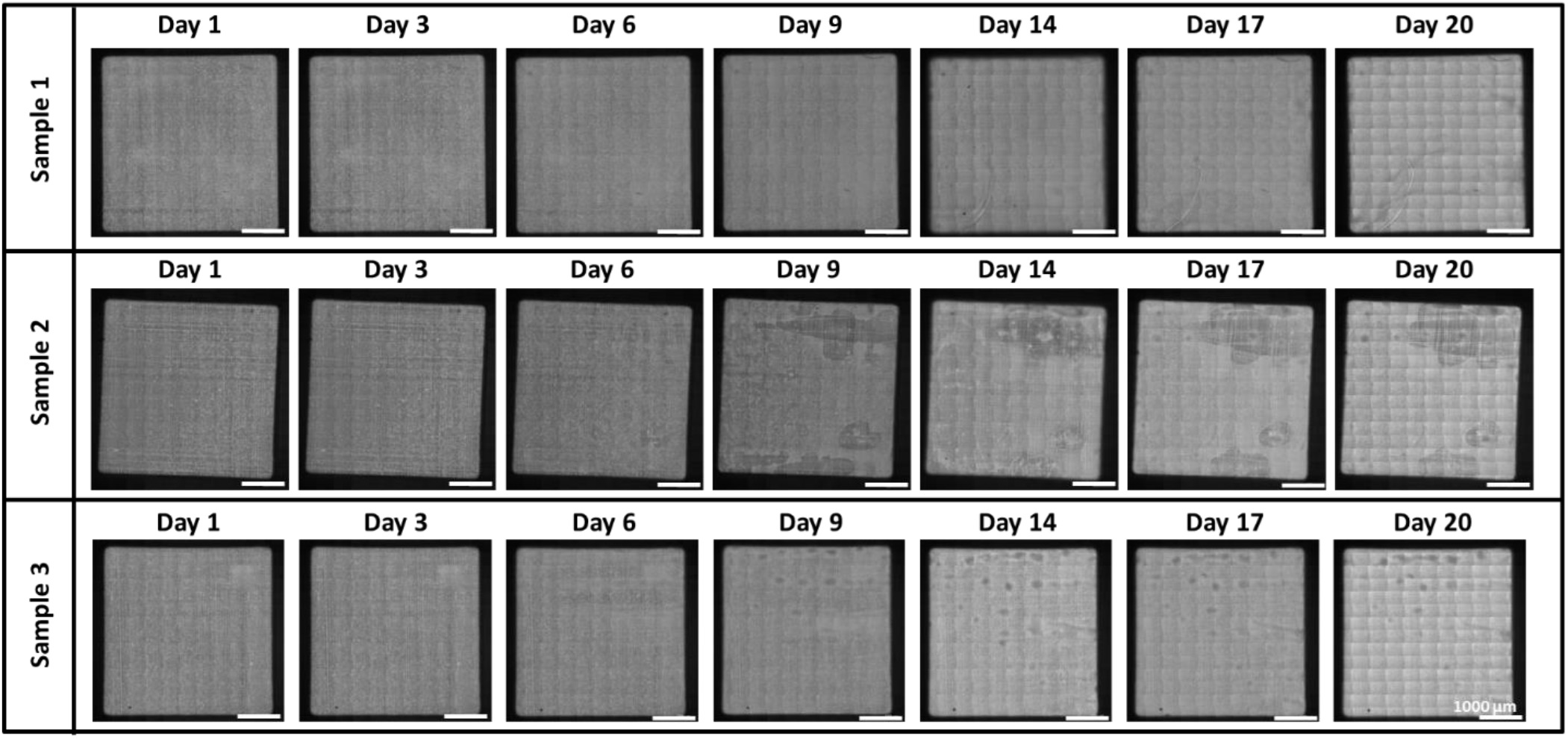
Mosaic images of pericyte cultures on crosshatch fiber networks with 800 nm fibers cultured for ∼3 weeks. (Scale bars: 1000 µm).

**Supplementary Figure 8:**
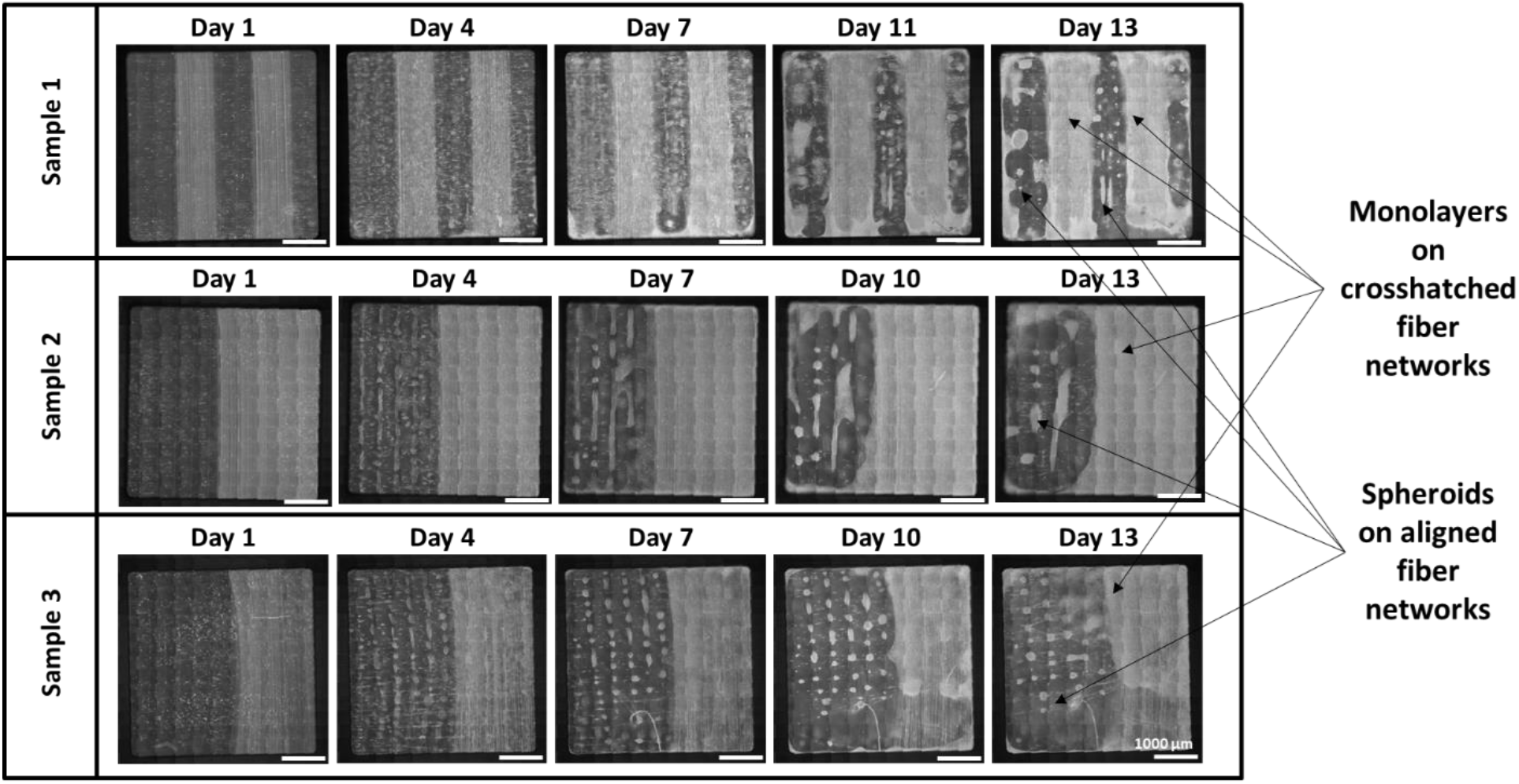
Mosaic images of pericyte cultures on fiber networks with aligned regions of 200 nm aligned fibers and 800 nm base fibers and crosshatch regions with 200 nm and 800 nm fibers deposited orthogonally cultured for ∼2 weeks. (Scale bars: 1000 µm).

**Supplementary Figure 9:**
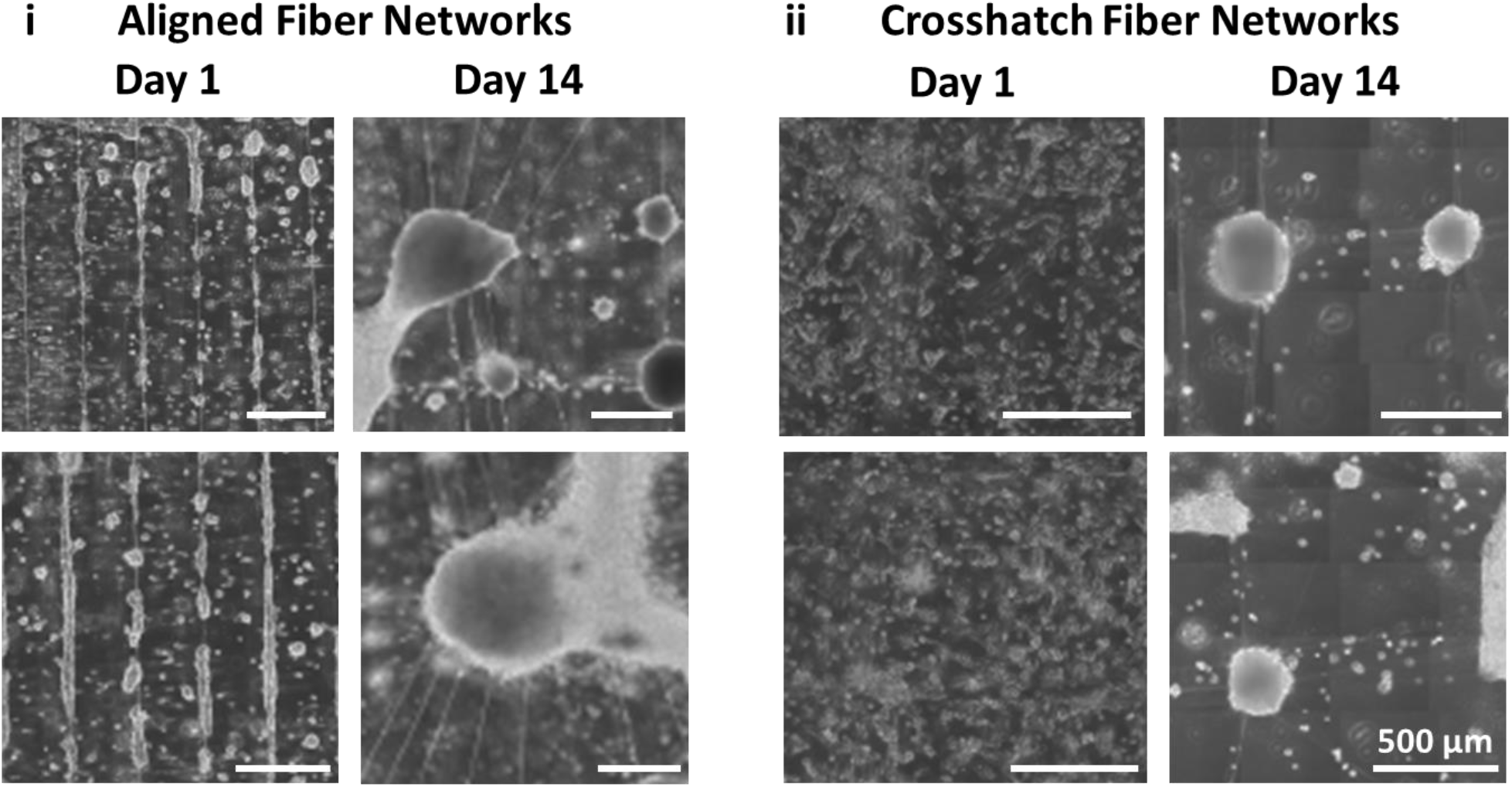
Spheroid formation involve extensive remodeling of the fiber networks in both (i) aligned architectures and (ii) crosshatched fiber networks. (Scale bars: 500 µm)

**Supplementary Figure 10:**
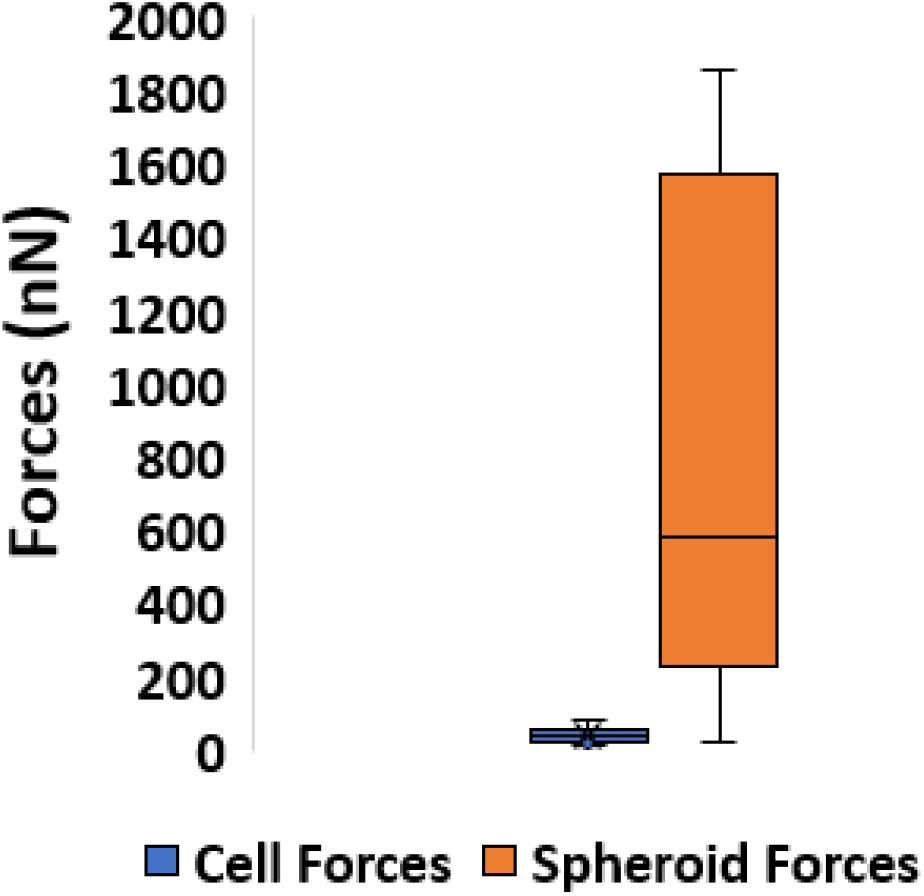
Comparison of cell forces (N=10) and spheroid forces (N=49)

**Supplementary Figure 11:**
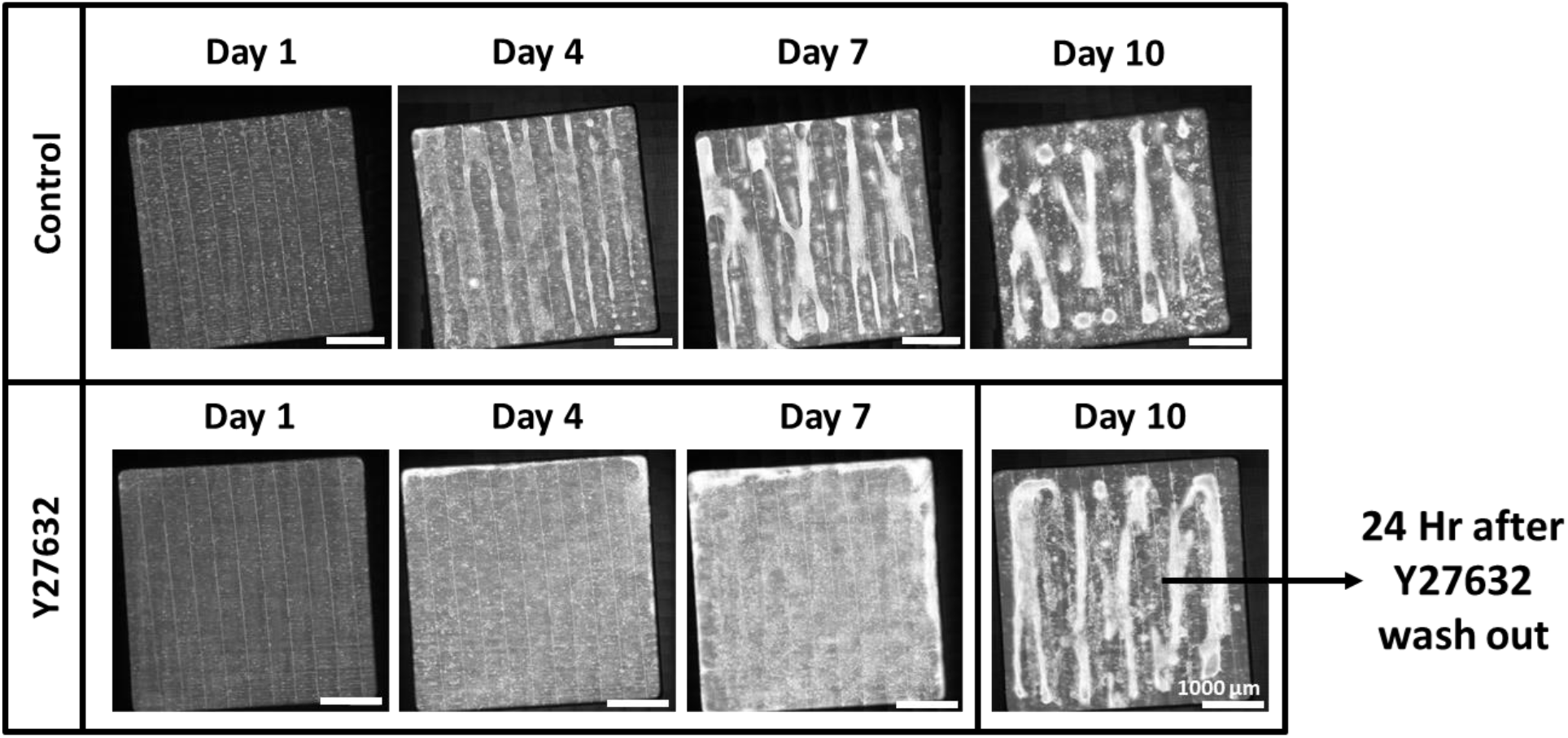
Growth of control pericyte cultures compared to pericyte cultures seeded with 20 µM Y27632 in the media. 24 hours after drug wash out on Day 10 show similar structures as seen in control cultures

## Movie Captions

Supplementary Movie 1: Multiple pericyte cells form a spheroid on aligned fiber networks by interacting with and contracting towards each other. (Scale bar: 50 μm)

Supplementary Movie 2: Multiple pericyte cells form a spheroid on aligned fiber networks by interacting with and contracting towards each other. (Scale bar: 50 μm)

Supplementary Movie 3: Multiple pericyte cells form a spheroid on aligned fiber networks by interacting with and contracting towards each other. (Scale bar: 50 μm)

Supplementary Movie 4: Elongated structures of aggregated pericytes contracting and remodeling the aligned fiber networks, breaking fused junctions and deforming the stiff 2 μm base fibers during the process (Scale bar: 50 μm).

Supplementary Movie 5: Elongated structures of aggregated pericytes contracting and remodeling the aligned fiber networks, breaking fused junctions and deforming the stiff 2 μm base fibers during the process (Scale bar: 50 μm).

Supplementary Movie 6: Elongated structures of aggregated pericytes contracting and remodeling the crosshatched fiber networks, breaking fused junctions and deforming the fibers during the process (Scale bar: 50 μm).

Supplementary Movie 7: Spheroid migrating along the aligned fibers towards the stiff 2 μm base fibers (Scale bar: 50 μm)

Supplementary Movie 8: A ‘leader’ cell pulling a spheroid along the 200 nm aligned fibers towards the stiff 2 μm base fibers (Scale bar: 50 μm)

Supplementary Movie 9: A ‘leader’ cell pulling a spheroid along the 200 nm aligned fibers (Scale bar: 50 μm)

Supplementary Movie 10: Spheroids on the stiff 2 μm base fibers in aligned fiber networks are predominantly stationary and use these fibers as a hub to grow in size (Scale bar: 50 μm)

Supplementary Movie 11: Spheroids on the stiff 2 μm base fibers in aligned fiber networks are predominantly stationary and use these fibers as a hub to grow in size (Scale bar: 50 μm)

Supplementary Movie 12: Spheroids on the stiff 2 μm base fibers in aligned fiber networks are predominantly stationary and use these fibers as a hub to grow in size (Scale bar: 50 μm)

Supplementary Movie 13: Spheroids continuously exchange cells with the surroundings over time, with cells moving into and sprouting out of the spheroid from multiple locations simultaneously (Scale bar: 50 μm).

Supplementary Movie 14: Spheroids continuously exchange cells with the surroundings over time, with cells moving into and sprouting out of the spheroid from multiple locations simultaneously (Scale bar: 50 μm).

Supplementary Movie 15: Spheroids continuously exchange cells with the surroundings over time, with cells moving into and sprouting out of the spheroid from multiple locations simultaneously (Scale bar: 50 μm).

Supplementary Movie 16: Two spheroids merge into a single larger spheroid over time when they come in contact with each other (Scale bar: 50 μm).

Supplementary Movie 17: Two spheroids merge into a single larger spheroid over time when they come in contact with each other (Scale bar: 50 μm).

Supplementary Movie 18: Two spheroids merge into a single larger spheroid over time when they come in contact with each other (Scale bar: 50 μm).

Supplementary Movie 19: Spheroids lose their contractility and hence shape when treated for 4 hours with 10 μM ROCK inhibitor Y27632. Upon washing off the drug and introduction of fresh culture media, spheroids are recovered within 12 hours (Scale bar: 50 μm).

Supplementary Movie 20: Spheroids lose their contractility and hence shape when treated for 4 hours with 10 μM ROCK inhibitor Y27632. Upon washing off the drug and introduction of fresh culture media, spheroids are recovered within 12 hours (Scale bar: 50 μm).

Supplementary Movie 21: Spheroids lose their contractility and hence shape when treated for 4 hours with 10 μM ROCK inhibitor Y27632. Upon washing off the drug and introduction of fresh culture media, spheroids are recovered within 12 hours (Scale bar: 50 μm).

Supplementary Movie 22: Spheroids lose their contractility and hence shape when treated for 8 hours with 20 μM ROCK inhibitor Y27632. Upon washing off the drug and introduction of fresh culture media, spheroids are recovered within 12 hours (Scale bar: 50 μm).

Supplementary Movie 23: Spheroids lose their contractility and hence shape when treated for 8 hours with 20 μM ROCK inhibitor Y27632. Upon washing off the drug and introduction of fresh culture media, spheroids are recovered within 12 hours (Scale bar: 50 μm).

Supplementary Movie 24: Spheroids lose their contractility and hence shape when treated for 15 hours with 20 μM ROCK inhibitor Y27632. (Scale bar: 50 μm).

Supplementary Movie 25: Spheroids lose their contractility and hence shape when treated for 15 hours with 20 μM ROCK inhibitor Y27632. (Scale bar: 50 μm).

Supplementary Movie 26: Spheroids lose their contractility and hence shape when treated for 15 hours with 20 μM ROCK inhibitor Y27632. (Scale bar: 50 μm).

Supplementary Movie 27: Pericytes seeded with 20 μM ROCK inhibitor Y27632 for a week grew into monolayers which when reversed, immediately (within 24 hrs) contracted towards each other and formed elongated cellular aggregates around the base fibers (Scale bar: 50 μm).

Supplementary Movie 28: Pericytes seeded with 20 μM ROCK inhibitor Y27632 for a week grew into monolayers which when reversed, immediately (within 24 hrs) contracted towards each other and formed elongated cellular aggregates around the base fibers (Scale bar: 50 μm).

Supplementary Movie 29: Pericytes seeded with 20 μM ROCK inhibitor Y27632 for a week grew into monolayers which when reversed, immediately (within 24 hrs) contracted towards each other and formed elongated cellular aggregates around the base fibers (Scale bar: 50 μm).

## References

1. Duval, K. et al. Modeling physiological events in 2D vs. 3D cell culture. Physiology vol. 32 266–277 (2017).

2. Baker, B. M. & Chen, C. S. Deconstructing the third dimension-how 3D culture microenvironments alter cellular cues. Journal of Cell Science vol. 125 3015–3024 (2012).

3. Achilli, T. M., Meyer, J. & Morgan, J. R. Advances in the formation, use and understanding of multi-cellular spheroids. Expert Opinion on Biological Therapy vol. 12 1347–1360 (2012).

4. Zanoni, M. et al. Modeling neoplastic disease with spheroids and organoids. Journal of Hematology and Oncology vol. 13 (2020).

5. Ong, C. S. et al. In vivo therapeutic applications of cell spheroids. Biotechnology Advances vol. 36 494–505 (2018).

6. Kim, S. jeong, Kim, E. M., Yamamoto, M., Park, H. & Shin, H. Engineering Multi-Cellular Spheroids for Tissue Engineering and Regenerative Medicine. Advanced Healthcare Materials vol. 9 (2020).

7. Shao, C. et al. Development of Cell Spheroids by Advanced Technologies. Advanced Materials Technologies vol. 5 2000183 (2020).

8. Mueller-Klieser, W. Multicellular spheroids - A review on cellular aggregates in cancer research. Journal of Cancer Research and Clinical Oncology vol. 113 101–122 (1987).

9. Sakalem, M. E., De Sibio, M. T., da Costa, F. A. da S. & de Oliveira, M. Historical evolution of spheroids and organoids, and possibilities of use in life sciences and medicine. Biotechnology Journal vol. 16 2000463 (2021).

10. Kelm, J. M., Timmins, N. E., Brown, C. J., Fussenegger, M. & Nielsen, L. K. Method for generation of homogeneous multicellular tumor spheroids applicable to a wide variety of cell types. Biotechnol. Bioeng. 83, 173–180 (2003).

11. Gerhardt, H. & Betsholtz, C. Endothelial-pericyte interactions in angiogenesis. Cell and Tissue Research vol. 314 15–23 (2003).

12. Armulik, A., Abramsson, A. & Betsholtz, C. Endothelial/pericyte interactions. Circulation Research vol. 97 512–523 (2005).

13. Dessalles, C. A., Babataheri, A. & Barakat, A. I. Pericyte mechanics and mechanobiology. Journal of Cell Science vol. 134 (2021).

14. Kumarasamy, M. & Sosnik, A. Heterocellular spheroids of the neurovascular blood-brain barrier as a platform for personalized nanoneuromedicine. iScience 24, 102183 (2021).

15. Cho, C. F. et al. Blood-brain-barrier spheroids as an in vitro screening platform for brain-penetrating agents. Nat. Commun. 8, 1–14 (2017).

16. Urich, E. et al. Multicellular self-assembled spheroidal model of the blood brain barrier. Sci. Rep. 3, 1–8 (2013).

17. Billaud, M. et al. Classification and Functional Characterization of Vasa Vasorum-Associated Perivascular Progenitor Cells in Human Aorta. Stem Cell Reports 9, 292–303 (2017).

18. Youssef, J., Nurse, A. K., Freund, L. B. & Morgan, J. R. Quantification of the forces driving self-assembly of three-dimensional microtissues. Proc. Natl. Acad. Sci. U. S. A. 108, 6993–6998 (2011).

19. Korff, T. & Augustin, H. G. Tensional forces in fibrillar extracellular matrices control directional capillary sprouting. J. Cell Sci. 112, 3249–3258 (1999).

20. Feng, F., Feng, X., Zhang, D., Li, Q. & Yao, L. Matrix Stiffness Induces Pericyte-Fibroblast Transition Through YAP Activation. Front. Pharmacol. 12, 1370 (2021).

21. Boot, R. C., Koenderink, G. H. & Boukany, P. E. Spheroid mechanics and implications for cell invasion. Advances in Physics: X vol. 6 (2021).

22. Efremov, Y. M. et al. Mechanical properties of cell sheets and spheroids: the link between single cells and complex tissues. Biophysical Reviews vol. 13 541–561 (2021).

23. Ahmadzadeh, H. et al. Modeling the two-way feedback between contractility and matrix realignment reveals a nonlinear mode of cancer cell invasion. Proc. Natl. Acad. Sci. U. S. A. 114, E1617–E1626 (2017).

24. Kopanska, K. S., Alcheikh, Y., Staneva, R., Vignjevic, D. & Betz, T. Tensile forces originating from cancer spheroids facilitate tumor invasion. PLoS One 11, (2016).

25. Nain, A. S., Sitti, M., Jacobson, A., Kowalewski, T. & Amon, C. Dry spinning based spinneret based tunable engineered parameters (STEP) technique for controlled and aligned deposition of polymeric nanofibers. Macromol. Rapid Commun. 30, 1406–1412 (2009).

26. Koons, B. et al. Cancer Protrusions on a Tightrope: Nanofiber Curvature Contrast Quantitates Single Protrusion Dynamics. ACS Nano 11, 12037–12048 (2017).

27. Padhi, A. et al. Force-exerting perpendicular lateral protrusions in fibroblastic cell contraction. Commun. Biol. 3, 1–11 (2020).

28. Singh, J., Pagulayan, A., Camley, B. A. & Nain, A. S. Rules of contact inhibition of locomotion for cells on suspended nanofibers. Proc. Natl. Acad. Sci. U. S. A. 118, (2021).

29. Jana, A. et al. Crosshatch nanofiber networks of tunable interfiber spacing induce plasticity in cell migration and cytoskeletal response. FASEB J. 33, 10618–10632 (2019).

30. Padhi, A. et al. Cell Fragment Formation, Migration, and Force Exertion on Extracellular Mimicking Fiber Nanonets. Adv. Biol. 5, 2000592 (2021).

31. Wintruba, K. L. et al. Adventitia-Derived Extracellular Matrix Hydrogel Enhances Contractility of Human Vasa Vasorum-Derived Pericytes via α2β1 integrin and TGFβ Receptor. J. Biomed. Mater. Res. - Part A (2022).

32. Sheets, K., Wang, J., Zhao, W., Kapania, R. & Nain, A. S. Nanonet Force Microscopy for Measuring Cell Forces. Biophys. J. 111, 197–207 (2016).

33. Hall, A. et al. Nanonet force microscopy for measuring forces in single smooth muscle cells of the human aorta. Mol. Biol. Cell 28, 1894–1900 (2017).

34. Padhi, A. & Nain, A. S. ECM in Differentiation: A Review of Matrix Structure, Composition and Mechanical Properties. Annals of Biomedical Engineering vol. 48 1071–1089 (2020).

35. Wei, Q. et al. Cellular modulation by the mechanical cues from biomaterials for tissue engineering. Biomater. Transl. 2, 323 (2021).

36. Smith, L. A. & Ma, P. X. Nano-fibrous scaffolds for tissue engineering. Colloids Surfaces B Biointerfaces 39, 125–131 (2004).

37. Wade, R. J. & Burdick, J. A. Engineering ECM signals into biomaterials. Materials Today vol. 15 454–459 (2012).

38. Ford, A. J. & Rajagopalan, P. Extracellular matrix remodeling in 3D: implications in tissue homeostasis and disease progression. Wiley Interdiscip. Rev. Nanomedicine Nanobiotechnology 10, e1503 (2018).

39. Van Helvert, S. & Friedl, P. Strain Stiffening of Fibrillar Collagen during Individual and Collective Cell Migration Identified by AFM Nanoindentation. ACS Applied Materials and Interfaces vol. 8 21946–21955 (2016).

40. Radisky, D., Muschler, J. & Bissell, M. J. Order and disorder: The role of extracellular matrix in epithelial cancer. in Cancer Investigation vol. 20 139–153 (2002).

41. Lu, P., Weaver, V. M. & Werb, Z. The extracellular matrix: A dynamic niche in cancer progression. Journal of Cell Biology vol. 196 395–406 (2012).

42. Taubenberger, A. V. et al. 3D extracellular matrix interactions modulate tumour cell growth, invasion and angiogenesis in engineered tumour microenvironments. Acta Biomater. 36, 73–85 (2016).

43. McKenzie, A. J. et al. The mechanical microenvironment regulates ovarian cancer cell morphology, migration, and spheroid disaggregation. Sci. Rep. 8, 1–20 (2018).

44. Jaiswal, D. et al. Stiffness analysis of 3D spheroids using microtweezers. PLoS One 12, (2017).

45. Lin, R. Z., Chou, L. F., Chien, C. C. M. & Chang, H. Y. Dynamic analysis of hepatoma spheroid formation: Roles of E-cadherin and β1-integrin. Cell Tissue Res. 324, 411–422 (2006).

46. Christiansen, J. J. & Rajasekaran, A. K. Reassessing epithelial to mesenchymal transition as a prerequisite for carcinoma invasion and metastasis. Cancer Research vol. 66 8319–8326 (2006).

47. Friedl, P. & Gilmour, D. Collective cell migration in morphogenesis, regeneration and cancer. Nature Reviews Molecular Cell Biology vol. 10 445–457 (2009).

48. Scarpa, E. & Mayor, R. Collective cell migration in development. J. Cell Biol. 212, 143–155 (2016).

49. Poujade, M. et al. Collective migration of an epithelial monolayer in response to a model wound. Proc. Natl. Acad. Sci. U. S. A. 104, 15988–15993 (2007).

50. Beaune, G. et al. Spontaneous migration of cellular aggregates from giant keratocytes to running spheroids. Proc. Natl. Acad. Sci. U. S. A. 115, 12926–12931 (2018).

51. Sharma, P. et al. Aligned fibers direct collective cell migration to engineer closing and nonclosing wound gaps. Mol. Biol. Cell 28, 2579–2588 (2017).

52. Haeger, A., Wolf, K., Zegers, M. M. & Friedl, P. Collective cell migration: Guidance principles and hierarchies. Trends in Cell Biology vol. 25 556–566 (2015).

53. Carey, S. P., Starchenko, A., McGregor, A. L. & Reinhart-King, C. A. Leading malignant cells initiate collective epithelial cell invasion in a three-dimensional heterotypic tumor spheroid model. Clin. Exp. Metastasis 30, 615–630 (2013).

54. Khalil, A. A. & Friedl, P. Determinants of leader cells in collective cell migration. Integrative Biology vol. 2 568–574 (2010).

55. Mayor, R. & Etienne-Manneville, S. The front and rear of collective cell migration. Nature Reviews Molecular Cell Biology vol. 17 97–109 (2016).

56. Lee, W. et al. Dispersible hydrogel force sensors reveal patterns of solid mechanical stress in multicellular spheroid cultures. Nat. Commun. 10, (2019).

57. Mark, C. et al. Collective forces of tumor spheroids in three-dimensional biopolymer networks. Elife 9, (2020).

58. Blumlein, A., Williams, N. & McManus, J. J. The mechanical properties of individual cell spheroids. Sci. Rep. 7, (2017).

59. Crosby, C. O. & Zoldan, J. Mimicking the physical cues of the ECM in angiogenic biomaterials. Regen. Biomater. 6, 61–73 (2019).

60. Tsai, A. C., Liu, Y., Yuan, X. & Ma, T. Compaction, fusion, and functional activation of three-dimensional human mesenchymal stem cell aggregate. Tissue Eng. - Part A 21, 1705–1719 (2015).

61. Devanny, A. J., Vancura, M. B. & Kaufman, L. J. Exploiting differential effects of actomyosin contractility to control cell sorting among breast cancer cells. Mol. Biol. Cell 32, ar24 (2021).

